# Spatial Decoupling of the Endodermal Functions Establishes the Innermost Cortex as the Gravity Sensor in Rice Stems

**DOI:** 10.64898/2026.10.31.763306

**Authors:** Rohan Kashyap, Zeenu Singh, Akshay Kumar, Yash Gupta, Amit Kumar, Shruti Singh, Stefan Jouanic, Véronique Bergounioux, Kalika Prasad, Pascal Gantet, Shri Ram Yadav

## Abstract

Gravitropism is a remarkable example of coordinated growth, enabling plants to perceive gravity and generate asymmetric cell expansion that restores organ orientation. In *Arabidopsis,* gravity sensing in shoots occurs within starch-rich endodermis cells. In rice, however, the stem must accommodate gravitropic bending while preserving integrity of tissues surrounding emerging crown roots, raising the possibility that gravity sensing relies on distinct structural and physiological adaptations. Here, using high-resolution imaging together with auxin-response analyses, we identify a specialized cell layer at the inner boundary of the rice stem cortex, operationally defined by its position adjacent to the lignified endodermis and associated with gravitropic responses. This layer is characterized by circumferentially elongated cells, starch sedimentation, a confined auxin-response domain, and expression of gravity-sensing markers. Following gravistimulation, starch granules realign and the auxin response domain expands asymmetrically within this layer, coinciding with differential growth during stem bending. Our results further indicate that this response depends on polar auxin transport and conserved components of the gravitropic regulatory network. Together, these findings support a model in which gravity sensing and signaling in rice stems are spatially separated from the lignified endodermal barrier. Such tissue organization may represent a developmental adaptation that allows the stem to reconcile barrier integrity with growth flexibility.

## Introduction

Plants are sessile organisms whose growth and responses to the environment rely on tightly coordinated patterns of cell division and expansion within tissues constrained by rigid cell walls. To support diverse physiological and developmental functions, specialized cell types are organized into distinct tissues and organs. A striking example is the endodermis. In roots, the endodermis forms a selective apoplastic barrier that seals the central vasculature and regulates the movement of water and solutes (*1–4*). In *Arabidopsis* shoots, the endodermis is commonly referred to as the starch-sheath (*5*). In several species, including *Arabidopsis*, starch-filled amyloplasts sediment within these cells in response to gravity, providing the mechanical cue that initiates gravitropic growth (*6–8*). As a result, barrier formation and gravity sensing have traditionally been viewed as complementary functions of homologous tissues in different organs. Whether this organizational framework is broadly conserved across flowering plants, however, remains an open question.

In monocot species such as rice (*Oryza sativa* L.), the stem serves as a central developmental hub, coordinating the formation of crown roots and tillers, and thereby shaping overall shoot and root systems architectures. The stem base, which is often submerged and is in direct contact with water, must function as a selective interface for water and nutrients uptake. Continuous emergence of post-embryonic crown roots therefore requires maintenance of an uninterrupted apoplastic barrier to prevent uncontrolled entry of water and solute into the vascular system. At the same time, the shoot must perceive and respond to gravity, an essential process for tiller angle determination and lodging resistance (*9–16*). These competing structural and physiological demands raise a fundamental question about tissue organization in species with a lignified stem endodermis as it is the case in rice (*17*). Lignification imparts the hydrophobic properties and mechanical strength required for endodermal barrier function (*8*, *18*), whereas gravitropic responses depend on highly dynamic processes including statolith sedimentation, auxin redistribution, and differential growth (*19–21*). The tissues in the rice stem base display dynamic changes of auxin response (*19–21*), however, it remained unclear whether auxin signaling is spatially organized within this developmental framework. In roots, these functions are spatially separated between the lignified endodermis and gravity-sensing columella cells (*19–21*). In rice seedlings, although amyloplast-sedimentation and presence of Casparian bands have been analyzed (*22–24*), how gravity perception and signaling is organized within the stem in the presence of a lignified endodermis remains poorly understood.

Here, we reveal a previously unrecognized auxin-signaling domain at the stem base of rice that lies immediately adjacent to the lignified endodermis. Based on its topological position overlying the endodermis, we designate this tissue as the innermost cortical cell layer (IC). Using confocal imaging, developmental analysis, physiological assays, data mining from available spatial transcriptomics (*25*) and RNA *in situ* hybridization, we show that the IC possesses hallmark features of gravity-sensing tissue, including localized auxin responses, statolith accumulation, and enrichment of gravitropism-associated genes. Gravistimulation induces dynamic, asymmetric auxin redistribution within this layer, while pharmacological and genetic analyses demonstrate contribution from polar auxin transport and conserved asymmetric growth regulators (*26*, *27*). Together, our findings support a model in which barrier function and gravitropic signaling are partitioned between adjacent cell layers in the rice stem base, revealing a developmental organization that reconciles the maintenance of a robust endodermal barrier with the cellular flexibility required for gravitropic growth.

## Results

### Endodermal Lignification Coincides with Auxin Signaling in the IC Layer

In rice and other cereals, post-embryonic development is characterized by the continuous emergence of crown roots, tiller buds, and leaves from the stem (Fig. 1A, Fig. S1) (*28*). Throughout this process the stem maintains a conserved radial organization in which the central stele is enclosed by a single-layered lignified endodermis that separates it from the surrounding cortex (Fig. 1B) (*23*). To identify potential sites of auxin response, we monitored the *DR5rev*::erGFP reporter activity (*29*, *30*). Stem sections were co-stained with Basic Fuchsin (BaFu) to visualise the lignified tissues and Fluorescent Brightener 28 (FB28) to reveal overall cell morphology (Fig. 1C‒E). Strikingly, we detected strong *DR5* reporter activity in vascular tissues and at the distal tips of crown root primordia (CRPs) (Fig. 1C‒C(ii)).

**Fig 1.**
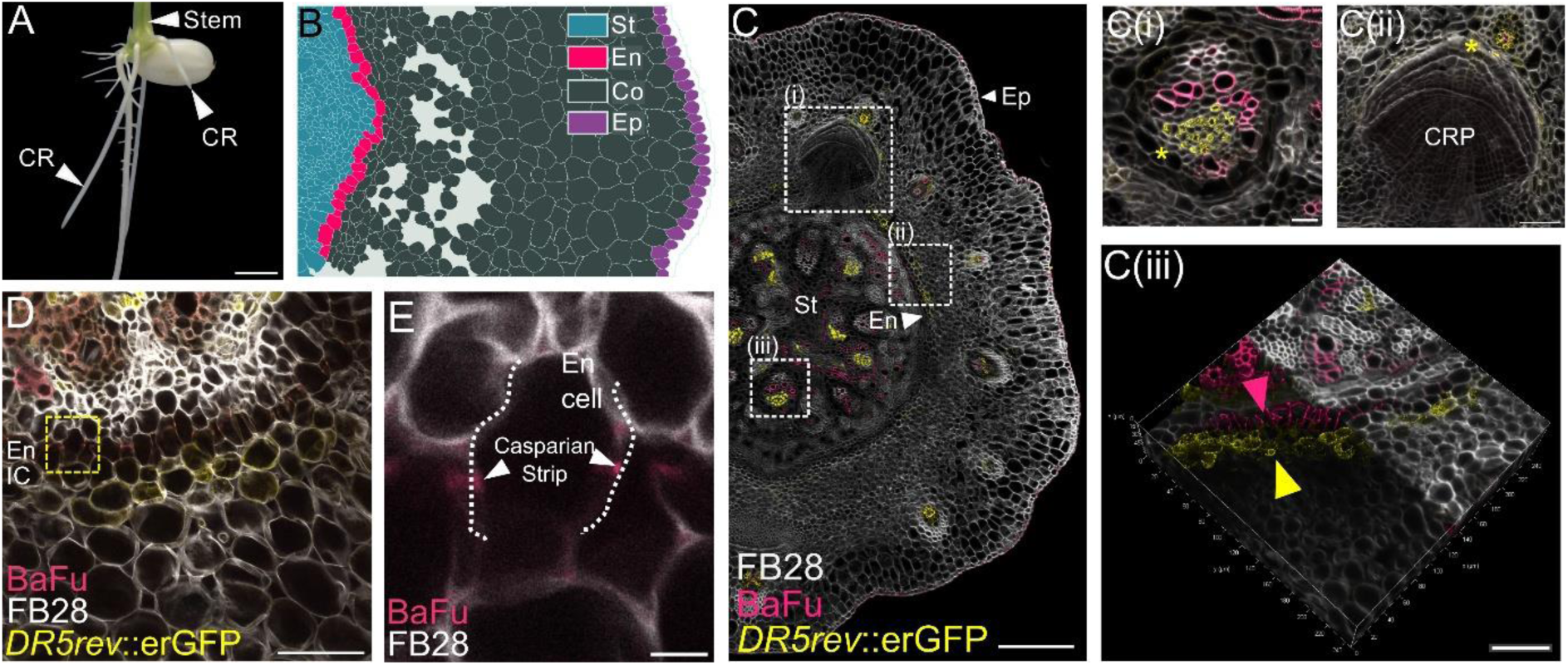
Endodermal lignification coincides with auxin signaling in the innermost cortical cell layer. *(A),* A 6-day-old rice seedling with crown roots (CR) originating from the stem, background removed for clarity (Scale bar: 0.2 cm). *(B),* An illustration showing the general anatomy of the rice stem with stele (St), endodermis (En), cortex (Co), and epidermis (Ep). *(C),* A section of the stem of a 6-day-old seedling with 3 major auxin signaling domains marked (*(i)* to *(iii)*; Scale bar: 200 μm). *(i)*, *DR5* signal in the stelar vasculature of the stem (yellow asterisk; Scale bar: 20 μm). *(ii)*, *DR5* signal at the tip of the developing crown root primordia (CRP, yellow asterisk; Scale bar: 50 μm). *(iii),* 3D projection of the lignified En (pink arrowhead) along with the adjacent *DR5* signal in the innermost cortical cell layer (yellow arrowhead; Scale bar: 50 μm). *(D)*, A Section of a 3-day-old stem base of *DR5rev*::erGFP lines with the En and the IC (Scale bar: 50 μm). *(E),* Dashed yellow square from *D*, an En cell showing the lignin deposition in a ‘casparian strip’ like fashion on the radial walls (dotted lines, Scale bar: 5 μm). FB28: Fluorescent Brightener 28, BaFu: Basic Fuchsin.

In addition, we observed a highly localized *DR5* signal in the cell layer situated immediately external to the lignified endodermis (Fig. 1C(iii)). Since this is the innermost layer of cortex adjacent to the endodermis, we refer to it as the innermost cortical cell layer (IC). Although endodermis-specific markers were not examined, the lignin deposition pattern on the radial cell walls was consistent with Casparian strip formation during early stages of endodermal lignification (Fig. 1D and *E*). As development progressed, these cells became extensively lignified beyond the Casparian strips (Fig. 1C(*iii*)). These anatomical features agree with previous descriptions of endodermal lignification in the mesocotyl and first internode of rice seedlings (*23*), supporting the topological assignment of this tissue. Together, these observations identify a previously unrecognized auxin response domain in the IC that emerges concurrently with endodermis lignification, suggesting that auxin responses are spatially organized immediately adjacent to the lignified endodermis of the rice stem.

### Auxin Signaling and Circumferential Growth Anisotropy Coincide in the IC layer

To determine how this spatial organization is established during development, we analyzed serial sections of 6-day-old stem bases and defined three developmental stages (Fig. 2 A‒C, Fig. S2). During Stage I, the endodermis was not stained by BaFu suggesting that it remained unlignified (Fig. 2A). Although CRP had already initiated at this stage, *DR5* activity was visibly undetectable in the IC cells (Fig 2A, Fig. S2*D*). In Stage II, the endodermis lignification, revealed by BaFu, coincided with the appearance of localized auxin response both at the distal tips of CRPs and within the IC layer (Fig. 2B, Fig. S2*D*). By Stage III, the lignified stem endodermis formed a continuous structural connection with the endodermis of the emerging crown roots, while *DR5* activity persisted in the adjacent IC cells (Fig 2C, Fig. S2*D*). To examine whether auxin perception follows a similar spatial pattern, we analyzed the DII-Venus auxin input reporter (Fig. S3) (*31*). During Stage I, the DII-Venus fluorescence was relatively uniform throughout the cortex. As development progressed through Stages II and III, however, DII-Venus fluorescence became stronger in the outer cortex, indicating lower auxin accumulation in these cells. This suggests although varying levels of auxin accumulates broadly across the cortex, auxin response is predominantly confined to the IC layer.

**Fig 2.**
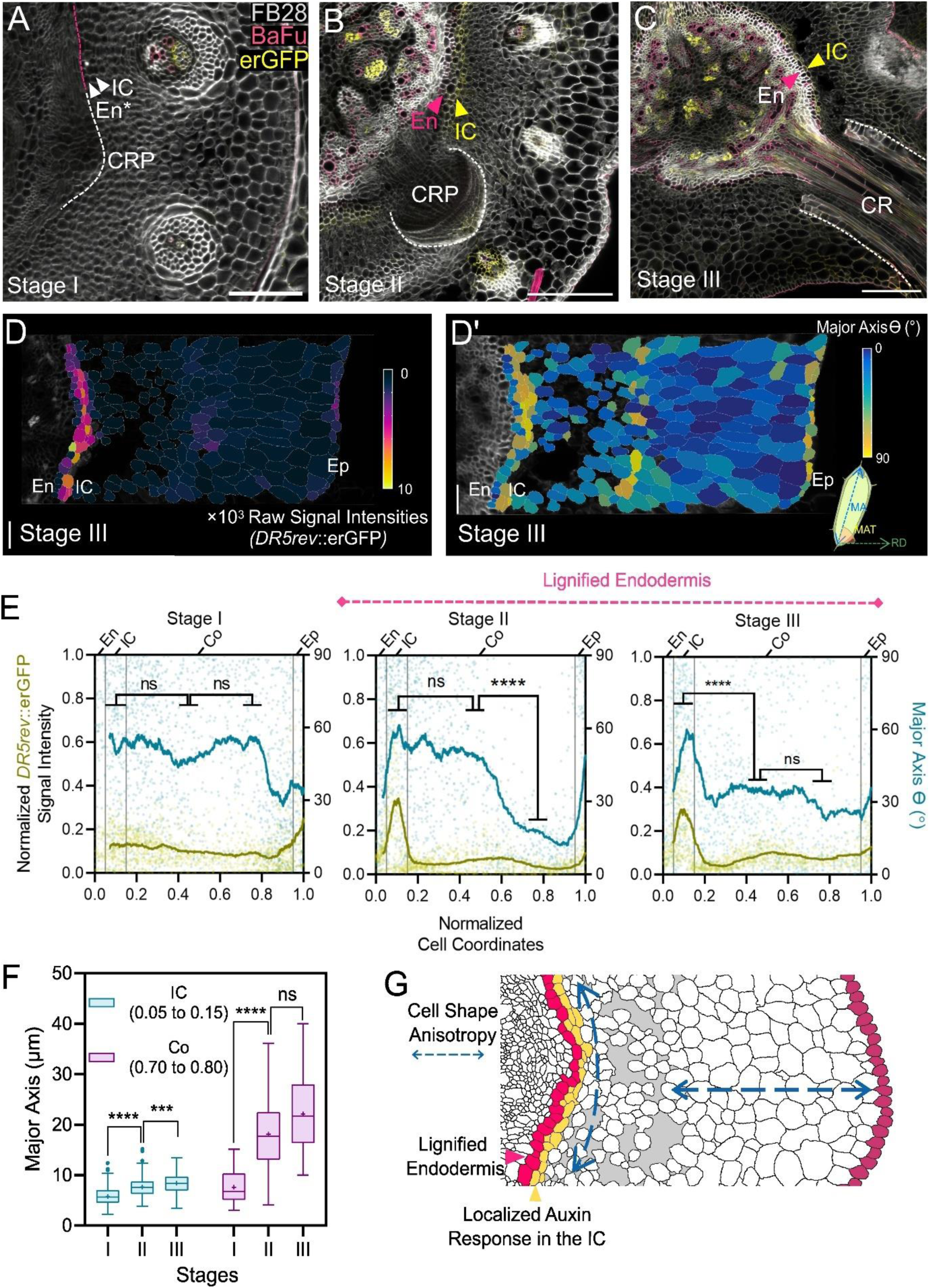
Auxin signaling and circumferential growth anisotropy overlap in the IC. *(A),* Representative section of a stem at stage I marked with initiating CRP (white-dotted line), unlignified endodermis (En*), and a lack of *DR5rev*::erGFP signal in the IC (Scale bar: 100 μm). *(B)*, Stem section at stage II with *DR5* signaling at the distal tip of the CRP, lignified En, and the appearance of the auxin response domain in the IC (Scale bar: 200 μm). *(C)*, Stem section at stage III, at which the En of the stem becomes continuous with the En of the CR, and the auxin signaling in the IC persists (Scale bar: 200 μm). *(D–D’),* Cellular segmentation of the rice stem bases at Stage III showing raw *DR5rev*::erGFP signal intensities *(D)* and the Major Axis Theta *(D’)* (MAθ, angle in degrees between the Major Axis, MA, of the cell and the reference direction, RD). The RD is measured along the stem radius (En: Endodermis, Ep: Epidermis; IC: Innermost Cortical Cell Layer, Scale bar: 50 μm). *(E)*, Graphical representation of normalized *DR5rev*::erGFP signal intensities (Left Y-axis) and MAθ (°, right Y-axis) of each cell plotted against normalized cell coordinates (0 = En and 1 = Ep, n = 6 for each stage, *\*\*\*\* p* < 0.0001 from Kruskal–Wallis test with Dunn’s multiple comparisons on the MAθ)*. (F),* Graphical representation of major axes of the cells of the IC (normalized cell coordinates between 0.05 and 0.15) and outer cortex (Co, 0.70 to 0.80) (*** *p* = 0.0003 and **** *p <* 0.0001 from Kruskal-Wallis test with Dunn’s multiple comparisons). *(G)*, An illustration showing the cell shape anisotropy of the cortical cells, and localized auxin response in the IC, adjacent to the lignified En.

Because cell morphology often reflects tissue specialization, we next quantified it using MorphoGraphX (*32*). We measured *DR5* signal intensity, along with major axis length and major axis theta (MAθ), which defines the orientation of cell elongation relative to radial axis of the stem (Fig. 2D–F, Fig. S4). Before endodermal lignification (Stage I), no GFP signal above the background level was detected across the cortex, and MAθ values showed substantial variation. With the onset of lignification in Stage II, both *DR5* activity and MAθ increased sharply within the IC layer. By Stage III, the MAθ peak became more pronounced, indicating that IC cells had adopted a circumferential growth orientation whereas cells in the outer cortex remained predominantly elongated along the radial axis. Despite this pronounced anisotropy, IC cells underwent only a modest increase in major axis length (46%), compared with 191% increase in the outer cortical cells (Fig. 2F). Together, our results show that the emergence of localized auxin response is tightly coupled to the acquisition of a distinctive circumferential cell geometry in the IC layer, defining this tissue as a morphologically and functionally specialized domain within the rice stem (Fig. 2G).

### The IC Layer Exhibits a ‘Statocyte-like’ Identity

In *Arabidopsis* hypocotyls, the unlignified endodermis forms a starch sheath in which starch-filled amyloplasts (statoliths) sediment to initiate gravity sensing and ultimately promote auxin redistribution to the adjacent tissues for asymmetric growth (*6–8*, *33*, *34*). In roots, by contrast, the heavily lignified endodermis functions primarily as an apoplastic barrier, whereas gravity sensing is delegated to the columella cells, which also exhibit strong *DR5* activity (*3*, *19–21*). We therefore hypothesized that rice stems similarly partition barrier and gravity-sensing functions between neighboring tissues. Previous studies have reported an endodermis/starch-sheath derived from the ground meristem at the rice stem base (*35*). Consistent with this, high-magnification bright-field imaging and Lugol staining revealed dense statolith accumulation in the IC cells, whereas the adjacent endodermis lacked detectable starch granules (Fig. 3A and *B*, Fig. S5).

**Fig 3.**
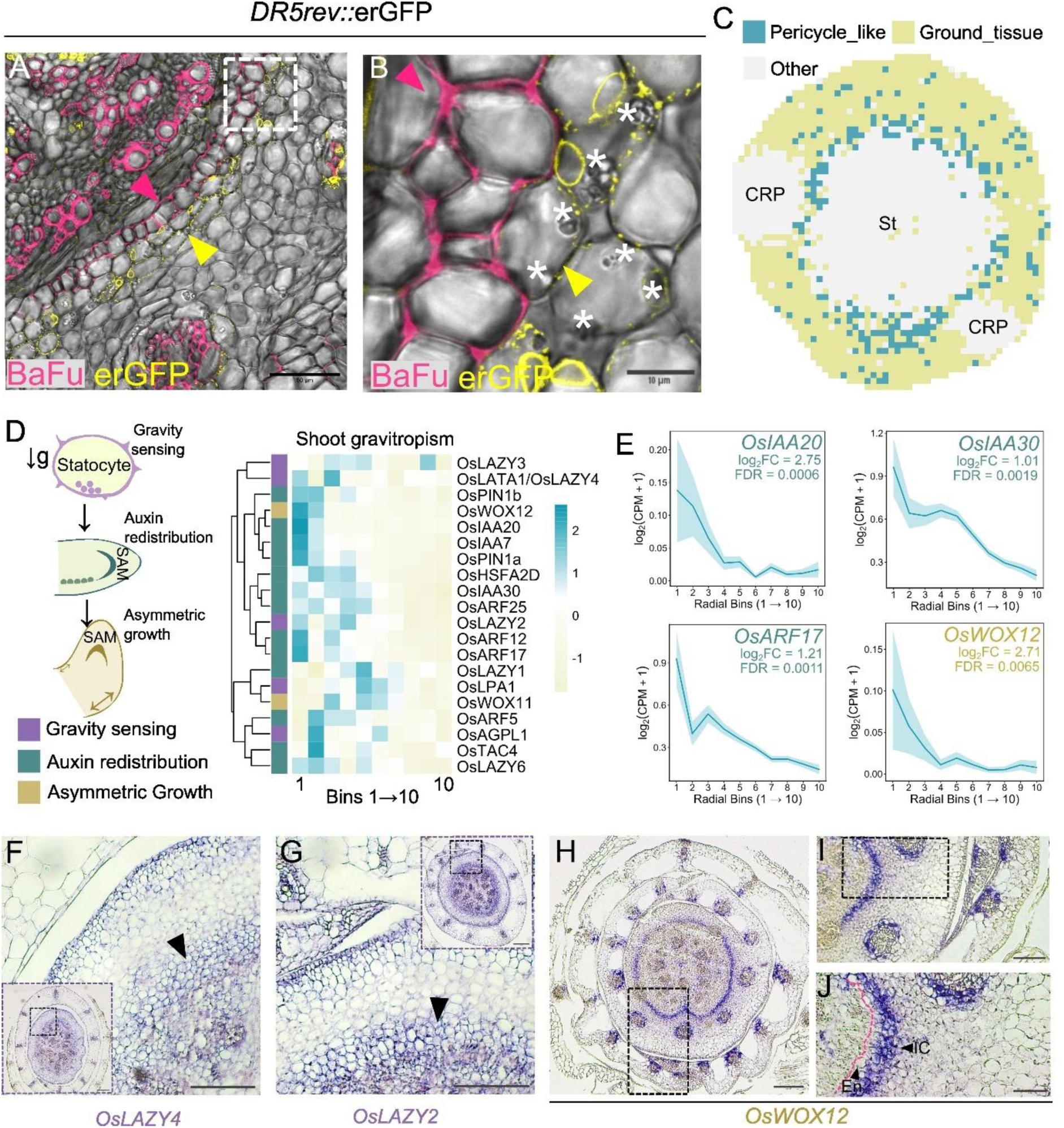
The Cells of the IC have a ‘statocyte-like’ identity. *(A–B)*, Sections of *DR5rev*::erGFP lines stained with Basic Fuchsin (BaFu) with trans-PMT channel, *(B)* is the cropped image from the dashed white square of *(A)*, note the occurrence of statoliths (white asterisks) in the cells that display the *DR5* signal. The pink and yellow arrowheads mark the En and the IC, respectively. *(C)*, The annotated tissue identities (‘Pericycle_like’ and ‘Ground_tissue’) on a spatial map of the rice stem base. The annotated cells were grouped into 10 radial bins, ordered from the innermost to the outermost relative to the centroid. The raw data used for this analysis were obtained from Zhong et al. (2025) (25), CRP: Crown Root Primordia, and St: Stele. *(D),* Heatmap showing the expression pattern of genes involved in shoot gravitropic response in shoots, divided into three processes: gravity sensing, auxin redistribution, and asymmetric growth (first column), displayed across all 10 radial bins. Expression values are shown as row-wise z-score calculated from log2(CPM+1). The genes were hierarchically clustered by rows. *(E),* Line plots, mean ± SEM log2(CPM+1) expression across radial bins (1 to 10) for representative genes. Log2FC and FDR were calculated by comparing bins 1 and 2 against all other bins. *(F–J)*, RNA *in situ* hybridization using antisense probes of *OsLAZY4 (F)*, *OsLAZY2 (G)*, *OsWOX12.1 (H–J)*, note the band-like expression pattern of the genes (black arrowhead) in *F* and *G*. *(I),* Dashed black rectangle from *H. (J),* Dashed black rectangle from *I*, note the specific expression pattern not only in the IC but also in the bundle sheath (Scale bar: 200 μm for *F* (inset)*, G* (inset), and *H*; 100 μm for *F, G,* and I; and 50 μm for *J*).

To further assess the “statocyte-like” identity of the IC layer, we analyzed the high-resolution spatial transcriptomic data from the rice stem base (*25*). Because the reference atlas did not resolve a distinct endodermal cluster, we used the annotated ‘Pericycle-like’, a layer just inner to the endodermis, and ‘Ground tissue’ cell populations as spatial approximations encompassing the stele-cortex boundary, the endodermis, and the surrounding ground tissue. Radial binning of these cells revealed significant enrichment of core gravitropism-associated genes in the innermost bins, which likely correspond to the IC layer (Fig. 3C–E, Table S1). These included the gravity-signaling components *OsLAZY6, OsLAZY2,* and *OsLAZY4* (*36*), together with auxin transporters (*OsPIN1a/b*), auxin signaling genes (*OsIAA20*, *OsIAA30*, and *OsARF17*), and *OsWOX12*, a regulator of cell expansion during gravitropic bending (27). We validated the expression of selected genes by RNA *in situ* hybridization. Notably, *OsLAZY2*, *OsLAZY4*, and *OsWOX12* showed a specific ring-like expression pattern surrounding the vasculature at the stem base (Fig. 3 F–H’, Fig. S6). The expression pattern of these 3 genes overlaps with the expression domain of *OsLAZY1,* a key regulator of polar auxin transport and shoot gravitropism in rice (*9*). Taken together, the topological position of IC layer adjacent to the lignified endodermis, combined with its localized auxin response, statolith accumulation, circumferential growth anisotropy, and enrichment of gravitropism-related genes, defines this layer as a distinct tissue with statocyte-like characteristics. Although definitive layer-specific transcript localization using cell-type-specific reporters will be required to confirm the precise cellular origin of these transcripts, the convergence of anatomical, physiological, and transcriptomic evidence strongly supports the IC as the principal gravity-associated signaling domain in the rice stem.

### Spatial Separation of Barrier Function and Gravitropic Signaling in the Rice Stem

To test whether the IC and the lignified endodermis perform distinct physiological functions, we next examined their respective roles in barrier formation and gravitropic signaling (Fig. 4). We first assessed the barrier function of the lignified endodermis using the *crown rootless1* (*crl1*) mutant, which lacks CRPs and therefore retains a continuous endodermis at the stem base (Fig. 4A, Fig. S7) (*37*). In a propidium iodide uptake assay, the dye was excluded by the lignified boundary, confirming that the endodermis functions as an effective apoplastic barrier independent of crown root emergence (Fig. 4B and *C*, Fig. S8).

**Fig 4.**
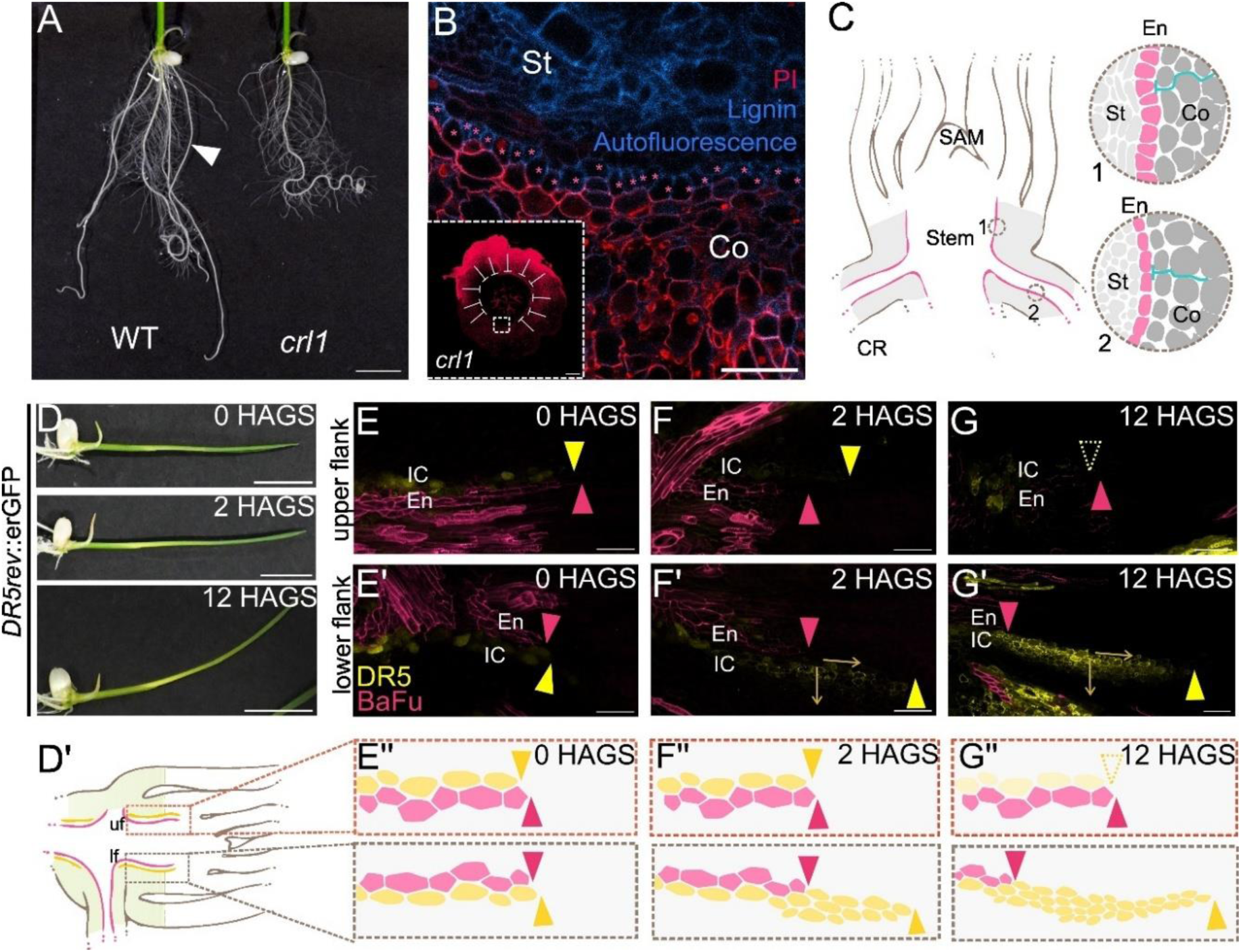
Spatial Separation of Stem Barrier Function and Gravitropic Signaling. *(A)*, Representative seedlings of 6-day-old wild-type (cv. Taichung 65) and *crl1* mutant, note the absence of crown roots in the mutant (white arrowhead in the WT, Scale bar: 1 cm). *(B)*, Propidium Iodide (PI) uptake assay in the stem of *crl1* (inset, scale bar: 200 μm), the dotted white square from the inset shows the restricted PI (represented by the white lines in the inset) at the lignified En (pink asterisks, marked with lignin autofluorescence, Scale bar: 50 μm). *(C)*, An illustration showing the continuity of the apoplastic barrier between the crown root (CR) and the stem. “1” and “2” mark the En of the Stem and CR, respectively. The blue line represents the blockage of the apoplastic path. *(D)*, 4-day-old seedlings at 0, 2, and 12 hours after gravistimulation (HAGS, Scale bar: 1 cm). *(E–G’)*, Upper (*E–G*) and lower flanks (*E’– G’*) of *DR5rev*::erGFP lines at 0 (*E* and *E’*), 2 (*F* and *F’*), and 12 (*G* and *G’*) HAGS. Note the spatial parity of lignin (pink arrowhead) and *DR5* signal (yellow arrowhead) in the upper flanks, and its eventual disappearance at 12 HAGS, contrasting with the expanding auxin signaling in the lower flanks (Scale bar: 50 μm). *(D’)*, Illustration of the longitudinal section of a stem base, marking the upper and lower flanks (uf and lf, respectively) used for Fig. *E– G’*. (*E’’–G’’*), Illustration showing the response of the IC layer HAGS, similar to Fig. *E–G’*.

We next investigated the role of the IC during gravity response using *DR5rev*::erGFP seedlings subjected to a 90° gravistimulation (Fig. 4D, and S10 *A–F*). Gravitropism in the stem involves gravity perception, asymmetric auxin distribution, and differential growth (*24*). Because the IC combines statoliths and a localized auxin response domain, we hypothesized that it contributes to the early stages of this process. Based on this, we first checked the sedimentation of the amyloplasts 1 hour after gravity stimulation (1 HAGS). We observed that the starch granules re-aligned along the direction of the gravity after reorienting seedlings 90° (Fig. S9). At 0 HAGS, the *DR5* activity in the IC closely followed the lignified endodermis boundary (Fig. 4E–E’’). Within 2 HAGS, well before visible stem bending occurred (Fig. 4*D*), the *DR5* activity on the lower flank extended apically beyond original alignment with the lignified endodermis (Fig. 4F–F’’). As the response progressed, this asymmetric auxin-response domain expanded further in both the apical and lateral directions across multiple cortical cell layers on the lower flank of the stem (Fig. 4 *G–G’’,* Fig. S10 *G–I’*). By 30 HAGS, the *DR5* activity had further extended upto the base of the coleoptile (the first leaf), a tissue previously implicated in gravity sensing in rice seedlings (Fig. S11*A* and *A’*) (*16*, *22*, *24*). Notably, a strong independent *DR5* signal was also evident in the lower flank of the coleoptile by 6 HAGS, suggesting a coordinated gravitropic response in stem and coleoptile (Fig. S11*B*).

To determine whether this asymmetric auxin response was accompanied by the changes in cellular organization, we quantified cell-shape at the coleoptile-stem junction using MorphoGraphX (dotted white square in Fig. S11*A*). We measured both *DR5* activity and the MAθ at the 30 HAGS. On the upper flank, the *DR5* peak became broader and less pronounced, whereas on the lower flank the auxin-response domain expanded basally, consistent with its asymmetric distribution (Fig. S11*C*). This spatial asymmetry was mirrored by changes in the cell shape. In the upper flank, MAθ peaked within the IC, while the cells in the remaining cortex averaged approximately 60°. In contrast, cells in the lower flank exhibited an average MAθ of approaching 80°, indicating enhanced circumferential growth anisotropy coincident with the expanding auxin response domain (Fig. S11*C*). Together, these physiological analyses provide functional support for a spatial separation of barrier function and gravitropic signaling within the rice stem. While the lignified endodermis retains its role as a robust apoplastic barrier, the adjacent IC exhibits the dynamic auxin redistribution and growth responses expected of a gravity-responsive tissue, supporting a model in which these complementary functions are partitioned between neighboring cell layers.

### Polar Auxin Transport and *OsWOX12* Coordinate Asymmetric Stem Growth after Gravistimulation

To investigate how the IC drives gravitropic responses, we combined pharmacological and genetic approaches. Because polar auxin transport is a central regulator of gravitropism in both shoots and roots (*20*), we first asked whether it is required for the asymmetric auxin response associated with the IC. Seedlings were treated with the auxin transport inhibitor 1-N-naphthylphthalamic acid (NPA) and *DR5* activity was monitored following gravistimulation. Compared with the DMSO control, NPA-treated seedlings showed a marked reduction in gravitropic bending that persisted throughout the 24 h observation period (Fig. 5A–B). By 24 HAGS, control seedlings had reached an average bending angle of approximately 60°, whereas NPA-treated seedlings bent only about 25° (Fig. 5B–C’). Consistent with this impaired response, the asymmetric *DR5* domain normally expanded along the lower flank in both the apical and lateral directions was largely abolished by NPA treatment (Fig. 5D and D’). These results indicate that polar auxin transport is required to establish the asymmetric auxin distribution that emerges from the IC during the early stages of gravitropism.

**Fig 5.**
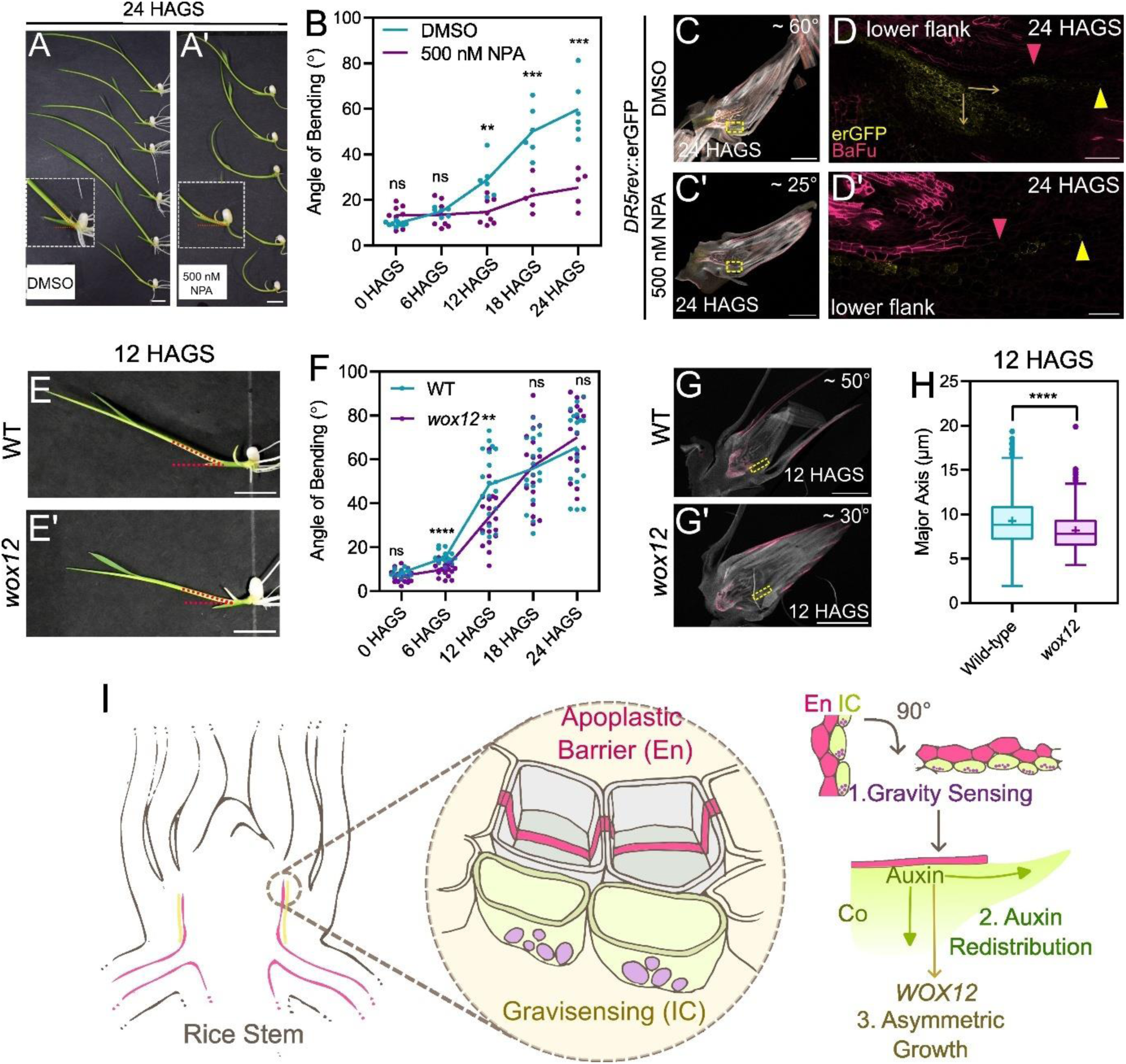
Polar Auxin Transport and *OsWOX12* Coordinate Asymmetric Stem Growth Downstream of the IC Domain. *(A–A’),* DMSO and 500 nM NPA-treated seedlings 24 hours after gravity stimulation (HAGS, Scale bar: 1 cm). *(B),* Graphical representation of the angle of bending (°) in DMSO vs NPA-treated seedlings at continuous HAGS (for DMSO: n=9 for 0 HAGS, and n=6 for the rest; for NPA: n=8 for 0, 6, and 12 HAGS; and n=5 for 18 and 24 HAGS; \*\**p* = 0.0023 for 12 HAGS, \*\*\**p =* 0.0008 for 18 HAGS, and *\*\*\*p* = 0.0006 for 24 HAGS from Student’s t test). *(C–C’),* Representative longitudinal section of DMSO (*C*) and NPA-treated (*C’*) seedlings 24 HAGS. The dashed-yellow square marks the region for Fig *D* and *D’* (Scale bar: 1 mm). *(D–D’),* Lower flanks of DMSO and NPA-treated seedlings 24 HAGS. Note the lack of expansion of the *DR5* signal in NPA treatment (Scale bar: 100 μm for DMSO and 50 μm for NPA). *(E–E’),* Wild-type (WT, *E*) and *wox12* (*E’*) seedlings 12 HAGS. Note the reduced bending in the *wox12* mutant (Scale bar: 1 cm). *(F),* Graphical representation of the angle of bending (°) in WT and *wox12* seedlings; n = 19; \*\*\*\**p* < 0.0001; and \*\**p* = 0.0039 from Student’s t test. *(G–G’)*, Representative longitudinal section of WT (*G*) and *wox12* (*G’*) seedlings 12 HAGS. The dashed-yellow square marks the region used for segmentation (Scale bar: 1 mm). *(H),* Graphical representation of major axes (µm) of the cortical cells from the lower flanks of WT and wox12; n > 400 from 3 independent seedlings, \*\*\*\**p* < 0.0001 from Mann-Whitney U test. *(I)*, An illustration summarizing the spatial decoupling of the endodermal functions within the En and the IC, along with the mechanism of action of the IC.

We next asked how this localized auxin response is translated into differential growth. Previous studies identified the transcription factors *OsWOX11* and *OsWOX6/12* as redundant regulators acting downstream of auxin signaling to promote asymmetric stem growth (*27*). To assess the contribution of this pathway, we examined the gravitropic response of the *wox12* mutant after transferring to dark conditions. Although the mutant seedlings displayed a clear delay in bending relative to wild-type during the first 12 HAGS, this defect gradually diminished, and by 24 HAGS the bending angles of the two genotypes were comparable (Fig. 5E–F). Cellular analysis of the lower flank further revealed that cells at the stem-coleoptile junction, where the *DR5* domain had expanded at 12 HAGS, displayed shorter major axes in the *wox12* mutant than in the wild type (Fig. 5G-H, Fig. S12). Together, these findings place the IC within a conserved gravitropic regulatory framework. Polar auxin transport is required to establish the asymmetric auxin-response domain associated with the IC, while *OsWOX12* contributes to the differential cell expansion that follows. These coordinated processes ultimately drive asymmetric stem growth and gravitropic bending. Together, these findings support a model in which IC contributes to gravitropic bending by engaging conserved components of the gravitropic pathway, particularly polar auxin transport, to drive asymmetric stem bending and suggest that this asymmetric stem bending is regulated at least in part by *OsWOX12* (Fig. 5I).

## Discussion

Gravity acts continuously on all plant cells, yet it is perceived only by specialized cells known as statocytes, which detect the direction of gravity and initiate downstream physiological responses (*38*). In angiosperms, starch-rich statocytes are typically located in the endodermis in shoots and the columella in roots, where sedimentable amyloplasts function as statoliths (*39*). An endodermis can inclusively be defined as a cell layer that has any two of the three criteria; (i) presence of Casparian strips, (ii) topologically the cell layer surrounding the vasculature, and (iii) expression of endodermis-specific genes such as SCR or its recognized ortholog (*3*). In *Arabidopsis*, the non-lignified starch-rich endodermis regulates shoot gravitropism and branch angle (*3*, *6*, *39*). In grasses, however, gravity perception has mainly been associated with the basal region of the coleoptile and leaf sheath (*16*, *22*, *24*). Here, we provide evidence that, in rice stems, gravity-associated signaling is concentrated in a distinct starch sheath layer, which we designate the inner cortex (IC), immediately adjacent to the lignified endodermis.

This finding revises the classical dual-function model in which the endodermis can act as an apoplastic barrier in root and the gravity-sensing tissue in shoot. Classical anatomical studies have noted that the walls of starch sheath of many stems remain unlignified (*17*). Our results support this distinction in rice stems, where the lignified endodermis appears specialized for barrier formation, the neighboring IC displays structural, transcriptional, and physiological characteristics consistent with a gravity-responsive tissue. Thus, barrier function and gravity sensing appear to be partitioned into adjacent cell layers, allowing the stem to maintain both apoplastic integrity and gravitropic responsiveness. A previous study reported that an actin-binding Rice Morphology Determinant (RMD) protein regulates light-dependent shoot gravitropism by regulating amyloplast sedimentation in the endodermal cells, however in absence of a complete anatomical tissue context or endodermis-associated marker, its identity as endodermis remained unestablished (*40*).

Current models propose that gravity perception begins with statolith sedimentation, followed by LAZY protein repolarization, PIN-mediated auxin redistribution, and differential cell expansion (*24*, *29*, *33*, *34*, *38–41*). These processes require dynamic intracellular organization and cellular flexibility, properties that may be incompatible with the rigid, lignified walls of the mature rice endodermis. In contrast, the IC provides a more permissive environment. IC cells sediment amyloplasts, express gravitropism-associated *LAZY* genes, display localized *DR5* activity, and show enhanced auxin response after gravistimulation. Although the lack of cell-type-specific markers and mutants prevents direct demonstration of gravity perception at single-cell resolution, the convergence of these anatomical, molecular, and physiological features strongly supports the IC as a statocyte-like tissue associated with gravitropic signaling.

The developmental sequence also supports this interpretation. CRPs already initiate from the innermost ground tissues of the stem during stage I, before extensive endodermal lignification occurs and IC-specific DR5 activity is established, however at stage II when CRPs progress for morphogenesis, the onset of auxin response coincides in IC and cap cells of the CRPs, raising a possibility that the IC and root cap tissues may share components of a common developmental program. Whether these tissues are developmentally related or evolutionarily homologous remains to be determined. This also raises yet another possibility whether the IC cells, in addition to gravity response, also contributes to root cap formation in the CRPs, similar to the case of lateral root primordia morphogenesis in a few species of Cucurbitaceae and Leguminosae families (*42*, *43*).

Although gravity-associated signaling appears to originate in the IC, the growth response likely involves coordinated activity across surrounding tissues. Following gravistimulation, the DR5 domain expands both apically and laterally before visible bending occurs. This differs from the predominantly lateral auxin redistribution reported in *Arabidopsis* hypocotyls, where PIN proteins redirect auxin laterally toward the epidermis (*33*, *34*). In rice, apical propagation may facilitate communication with younger, more extensible tissues near the stem apex and coleoptile base, coordinating differential growth across the organ. Consistent with this idea, later stages of bending were accompanied by asymmetric cell elongation and differentiation of new lignified tissues on the responding flank. However, because tissue mechanics were not measured directly, the relationship between auxin redistribution and differential growth remains inferential.

Our findings also extend current models of gravity sensing in grasses. The bases of coleoptile and leaf sheath have been regarded as the principal gravitropic organ in rice seedlings (*16*, *22*, *24*), whereas we now report the IC for gravity response in the rice stem. Previous studies showed that *OsLAZY2* and *OsLAZY3* regulate shoot gravitropism and tiller angle through a starch-statolith pathway in leaf sheath base cells, whereas *OsLAZY4* acts through a parallel pathway. Both converge upstream of *OsLAZY1* to establish asymmetric auxin distribution and activate *OsWOX6/12* and *OsWOX11* (*16*, *27*, *44*, *45*). Rather than replacing this model, our results suggest that the rice stem IC is an additional site of the same gravitropic network. During gravistimulation, auxin response expanded from the stem IC laterally toward the coleoptile base, indicating that these tissues function together as an integrated gravity-sensing system. Our spatial expression pattern analysis reveals that though the gravity-sensing genes, *OsLAZY2* and *OsLAZY4* express in the multiple cortical layers, similar to *OsLAZY1* (*9*), the auxin response and its downstream effector (*OsWOX12*) are activated in a more confined IC layer.

The ontological origin and molecular basis of IC specification remains unknown. In *Arabidopsis*, the conserved SHORT-ROOT (SHR) pathway specifies the shoot starch sheath endodermis (*6*, *46*), whereas in monocot roots, including maize and *Setaria*, SHR protein movement extends across multiple cortical layers (*47*). It is therefore plausible that related developmental mechanisms specify both the endodermis and IC in rice stems, partitioning endodermal functions between neighboring tissues. Testing this hypothesis will require direct genetic and cell-biological analyses and should clarify the developmental identity of the IC.

Our findings may also have evolutionary implications. Early monocot lineages are thought to have diversified in aquatic or semi-aquatic environments (*48–50*). In such conditions, increased endodermal lignification may have strengthened the apoplastic barrier while limiting its capacity to support dynamic gravitropic bending. The evolution of a neighboring starch sheath, represented here by the IC, could therefore have preserved gravity sensing without compromising barrier function. Although speculative, this scenario provides a framework for understanding how functional specialization of stem tissues may have diverged between monocots and eudicots. Finally, these findings have practical significance for crop architecture. Shoot gravitropism is a major determinant of tiller angle, a trait that strongly influences planting density, canopy architecture, and grain yield in rice (*9–12*, *16*, *26*, *27*, *51*). Identifying the IC as a potential site of gravity-associated signaling expands the current framework of rice shoot gravitropism and provides a foundation for future studies aimed at manipulating cereal architecture through tissue-specific regulation of gravity responses.

## Materials and Methods

### Plant material and growth conditions

The *DR5rev*::erGFP and *wox12* lines used in the study are in *Oryza sativa* L. (cv. Taipei309). Mutant for *wox12* was generated using CRISPR-Cas9, two guide RNAs were designed in the first exon and were cloned in the *p*RGEB32 vector following Xie et al. 2015 (*52*). The construct was transformed into *Agrobacterium tumefaciens* LBA4404, and was used for plant transformation as described previously (*30*, *53*). The *crl1* mutant line is in the cv. Taichung 65 background. Seeds were surface-sterilized with 70% ethanol for 90 seconds and 40% (v/v) Rin-ALA (commercial bleach) for 20 minutes, followed by repeated washes with sterile water. The seeds were soaked in water for 24 hours and then placed in ½ Murashige and Skoog (MS) media (TS1004, Himedia) containing 1% (w/v) sucrose (Himedia) and 0.05% (w/v) MES (Himedia), solidified with 0.35% (w/v) phytagel (Sigma), and adjusted to pH 5.8. Media was poured into glass jam bottles or magenta boxes. Seedlings were grown at 28°C under a 16 h light:8 h dark photoperiod. For NPA treatment, a 1 mM stock of Naptalam (Sigma) was prepared in DMSO, filtered-sterilized, and added to the autoclaved media before pouring.

### Vibratome sectioning and confocal microscopy

All tissues were fixed in 4% paraformaldehyde (Sigma) in 1× PBS overnight at 4°C, after 20 minutes of vacuum infiltration. The tissues were then washed once with 1× PBS and embedded in 4% agarose (SeaKem) for vibratome sectioning using a Leica VT1000S. Sections of 70-80 μm were transferred to the ClearSee solution for 2 days (*54*). The sections were subsequently dual-stained with 0.1% Calcofluor White (Himedia) and 0.2% Basic Fuchsin (Himedia) prepared in ClearSee (*55*), and mounted in ClearSee for confocal microscopy. Imaging was performed using a Leica Stellaris 5 microscope with 63×/1.40 oil–immersion and 20×/0.75 dry objectives. Image acquisition parameters were set to 1 Airy unit, a 1024×1024-pixel frame size, line averaging of 2, and a bidirectional scan speed of 400 Hz. Due to the large tissue sizes, all macroscopic images were acquired as tile scans. Fluorescent Brightener 28 (FB28) was excited with a 405 nm laser at 0.87% power (gain: 2.5; emission range: 425–483 nm. Basic Fuchsin was excited with a 561 nm laser at 10.86% power (gain: 23.4; emission range: 600–650 nm). erGFP was excited with a 488 nm laser at 12.31% power (gain: 26; emission range: 503–535 nm). The images were scaled and processed using Fiji software (https://imagej.net/) and Leica LasX software (*56*).

### MorphoGraphX analysis

The TIFF files exported from Fiji were used for the MorphoGraphX (MGX) analysis (*32*). Briefly, imported stacks were blurred with a Gaussian kernel and used for edge detection at a threshold of 1. The flat plane of the stack was used to create an overlying mesh, which was segmented and smoothed to a target size of 500 nm. The signal from the FB28 channel was projected onto the mesh from a single z-slice. Cells were manually seeded, followed by watershed segmentation. All the segmentations were manually checked and corrected for any errors. Major axis theta and signal intensities were quantified using the heatmap function after projection of the respective channels. The values were then exported as CSV files for plotting.

### Gravity stimulation experiment

Seedlings were grown in magenta boxes as described above. Four days post-sterilization, the boxes were rotated 90° and transferred to the dark. Seedlings were imaged every 6 hours, and corresponding tissues were collected for sectioning. The angle of bending was quantified using the angle tool Fiji.

### Propidium iodide uptake experiment

*crl1* mutant (cv. Taichung65) seedlings were grown as described above. The stem bases of 6-day-old seedlings were collected and immersed in a 15 μg.mL^-1^ solution of propidium iodide (PI, Sigma) in water. The stems were vacuum-infiltrated with the PI solution for 20 minutes and incubated for 1 hour at 30°C. Tissues were then washed with water and embedded in 4% agarose. Sections of 200 μm were cut using a Leica VT1000S vibratome and mounted in water. Imaging was performed using a 20x/0.75 DRY objective, followed by a 63x/1.40 oil-immersion objective. Image acquisition parameters were set to 1 Airy unit, a 1024x1024 pixel frame size, line averaging 2, and a scan speed of 400 Hz, with bidirectional scanning. PI was excited with a 514 nm laser at 2% power (gain: 26.4; emission range: 550–740 nm). Lignin autofluorescence was captured using a 405 nm laser at 15% power (gain: 51.3; emission range: 425–469 nm).

### Spatial transcriptomics data re-analysis

To analyze the spatial expression patterns of the canonical gravitropic regulatory modules, raw transcriptomic data and corresponding cell-type annotations were retrieved from STOmics DB (http://db.cngb.org/stomics) under the STT0000026 identifier (*25*, *57*). The spatial analysis was restricted to the cells annotated as “Pericycle_like” and “Ground_tissue”. To establish a standardized radial axis across varying stem cross–sections, the spatial coordinates for each sample were scaled, and the developmental center of the stem base was defined by calculating a spatial centroid of the pericycle cell population. The Euclidean distance of each cell from this centroid was calculated to determine its absolute radial position. These radial distances were min-max normalized and were partitioned into 10 equally spaced bins. A set of 20 locus IDs (Table S1) associated with the gravitropic response was used for the analysis. Raw transcript counts (MIDCounts) were normalized to counts per million (CPM). A subsequent log2(CPM+1) transformation was applied to stabilize variance across the dataset. For each spatial bin, log2(CPM+1) expression values were averaged per sample, followed by the calculation of a global mean expression profile per spatial bin. To generate the heatmap using the pheatmap R package, the global mean expression for each gene was standardized into row-wise Z-scores across the 10 radial bins. For individual gene trajectories, the mean expression and standard error of the mean (±SEM) were calculated across all cells within each bin and visualized using the ggplot2 package. To quantify the magnitude of spatial segregation within the inner cortex (Bins 1 and 2) relative to the outer (Bins 3–10), a linear fold-change was calculated as the ratio of inner and outer expression values, which was log2-transformed to generate the final reported *log2*FC values. To assess the statistical significance of these spatial differences, raw p-values were calculated using Fisher’s exact tests to compare the proportion of expressing versus non–expressing cells between the inner and the outer radial bins. These p-values were corrected for multiple hypothesis testing using the Benjamini-Hochberg False Discovery Rate (FDR). R version 4.4.1 was used for all the statistical analyses (https://www.R-project.org/).

### RNA in situ hybridization

Gene-specific fragments of *OsLAZY2, OsLAZY4,* and *OsWOX12* were cloned into pBluescript SK+, linearized with HindIII, and transcribed with T3 RNA Polymerase (Roche) for DIG-labeled antisense probes. EcoRI linearized clones were used to synthesize the sense probes with T7 RNA Polymerase (Roche). Stem bases of 6-day-old seedlings were fixed in 4% paraformaldehyde (Sigma-Aldrich) in 1X PBS overnight. The tissues were dehydrated using a graded ethanol series (30%, 50%, 70%, and 85%) with 130 mM NaCl (Sigma-Aldrich). This was followed by a final incubation with 95% and 100% ethanol. 10 μm-thick sections were taken on poly-L-lysine-coated slides and were hybridized as described previously. Briefly, the slides were dewaxed with xylene (Himedia), hydrated in an ethanol series, acidified, and treated with 1 μg mL^-1^ proteinase K (Sigma-Aldrich). After re-fixing in 4% paraformaldehyde, the slides were hybridized with DIG-labeled probes overnight at 55°C in hybridization chambers, soaked in 50% formaldehyde solution (Rankem). The slides were then washed and treated with 10 μg mL^-1^ RNase (Himedia) to degrade unbound probes. 0.5% Boehringer Mannheim reagent was used for blocking, prior to anti-DIG antibody (Sigma-Aldrich) treatment. The signal was detected using NBT-BCIP (Sigma-Aldrich). Dehydrated slides were then mounted in Entellan (Sigma-Aldrich) and were imaged using a Leica Visoria B microscope with 4X and 10X objectives.

## Acknowledgments

The Department of Biotechnology (DBT), Government of India, the Anusandhan National Research Foundation (ANRF), Government of India, and Indo-French Centre for the Promotion of Advanced Research (IFCPAR/CEFIPRA)/Department of Science and Technology, Government of India, are acknowledged for financial support to the projects of S.R.Y.’s laboratory. Fellowships are provided by the Council of Scientific & Industrial Research (CSIR), India, to R.K., by the Department of Biotechnology (DBT), India, to S.S., and Amit K. and by IIT Roorkee to Y.G., and A.K. We acknowledge the Ashok Soota Molecular Medicine Facility at the Department of Biosciences and Bioengineering, IIT Roorkee, for access to the confocal microscope.

## References

1. J. A. Raven, D. Edwards, Roots: evolutionary origins and biogeochemical significance. Journal of Experimental Botany 52, 381–401 (2001).

2. S. Naseer, Y. Lee, C. Lapierre, R. Franke, C. Nawrath, N. Geldner, Casparian strip diffusion barrier in Arabidopsis is made of a lignin polymer without suberin. Proc Natl Acad Sci U S A 109, 10101–10106 (2012).

3. N. Geldner, The Endodermis. Annu. Rev. Plant Biol. 64, 531–558 (2013).

4. M. Barberon, The endodermis as a checkpoint for nutrients. New Phytologist 213, 1604–1610 (2017).

5. A. M. F. Tomescu, The stele – a developmental perspective on the diversity and evolution of primary vascular architecture. Biol Rev 96, 1263–1283 (2021).

6. H. Fukaki, J. Wysocka-Diller, T. Kato, H. Fujisawa, P. N. Benfey, M. Tasaka, Genetic evidence that the endodermis is essential for shoot gravitropism in *Arabidopsis thaliana*. The Plant Journal 14, 425–430 (1998).

7. K. Fujihira, T. Kurata, M. K. Watahiki, I. Karahara, K. T. Yamamoto, An Agravitropic Mutant of Arabidopsis, endodermal-amyloplast less 1, That Lacks Amyloplasts in Hypocotyl Endodermal Cell Layer. Plant and Cell Physiology 41, 1193–1199 (2000).

8. M. T. Morita, T. Kato, K. Nagafusa, C. Saito, T. Ueda, A. Nakano, M. Tasaka, Involvement of the Vacuoles of the Endodermis in the Early Process of Shoot Gravitropism in Arabidopsis. Plant Cell 14, 47–56 (2002).

9. P. Li, Y. Wang, Q. Qian, Z. Fu, M. Wang, D. Zeng, B. Li, X. Wang, J. Li, LAZY1 controls rice shoot gravitropism through regulating polar auxin transport. Cell Res 17, 402–410 (2007).

10. T. Yoshihara, M. Iino, Identification of the Gravitropism-Related Rice Gene LAZY1 and Elucidation of LAZY1-Dependent and -Independent Gravity Signaling Pathways. Plant and Cell Physiology 48, 678–688 (2007).

11. X. Wu, D. Tang, M. Li, K. Wang, Z. Cheng, Loose Plant Architecture1, an INDETERMINATE DOMAIN Protein Involved in Shoot Gravitropism, Regulates Plant Architecture in Rice. Plant Physiology 161, 317–329 (2012).

12. M. Okamura, T. Hirose, Y. Hashida, T. Yamagishi, R. Ohsugi, N. Aoki, Starch reduction in rice stems due to a lack of OsAGPL1 or OsAPL3 decreases grain yield under low irradiance during ripening and modifies plant architecture. Functional Plant Biology 40, 1137–1146 (2013).

13. Z. Li, Y. Liang, Y. Yuan, L. Wang, X. Meng, G. Xiong, J. Zhou, Y. Cai, N. Han, L. Hua, G. Liu, J. Li, Y. Wang, OsBRXL4 Regulates Shoot Gravitropism and Rice Tiller Angle through Affecting LAZY1 Nuclear Localization. Molecular Plant 12, 1143–1156 (2019).

14. Y. Hu, S. Li, X. Fan, S. Song, X. Zhou, X. Weng, J. Xiao, X. Li, L. Xiong, A. You, Y. Xing, *OsHOX1* and *OsHOX28* Redundantly Shape Rice Tiller Angle by Reducing *HSFA2D* Expression and Auxin Content. Plant Physiol. 184, 1424–1437 (2020).

15. H. Li, H. Sun, J. Jiang, X. Sun, L. Tan, C. Sun, *TAC4* controls tiller angle by regulating the endogenous auxin content and distribution in rice. Plant Biotechnology Journal 19, 64–73 (2021).

16. L. Huang, W. Wang, N. Zhang, Y. Cai, Y. Liang, X. Meng, Y. Yuan, J. Li, D. Wu, Y. Wang, *LAZY2* controls rice tiller angle through regulating starch biosynthesis in gravity-sensing cells. New Phytologist 231, 1073–1087 (2021).

17. J. L. Seago, Revisiting the occurrence and evidence of endodermis in angiosperm shoots. Flora 273, 151709 (2020).

18. L. J. Gibson, The hierarchical structure and mechanics of plant materials. J. R. Soc. Interface. 9, 2749–2766 (2012).

19. S. Sabatini, D. Beis, H. Wolkenfelt, J. Murfett, T. Guilfoyle, J. Malamy, P. Benfey, O. Leyser, N. Bechtold, P. Weisbeek, B. Scheres, An Auxin-Dependent Distal Organizer of Pattern and Polarity in the Arabidopsis Root. Cell 99, 463–472 (1999).

20. J. Friml, J. Wiśniewska, E. Benková, K. Mendgen, K. Palme, Lateral relocation of auxin efflux regulator PIN3 mediates tropism in Arabidopsis. Nature 415, 806–809 (2002).

21. M. Taniguchi, M. Furutani, T. Nishimura, M. Nakamura, T. Fushita, K. Iijima, K. Baba, H. Tanaka, M. Toyota, M. Tasaka, M. T. Morita, The Arabidopsis LAZY1 Family Plays a Key Role in Gravity Signaling within Statocytes and in Branch Angle Control of Roots and Shoots. Plant Cell 29, 1984–1999 (2017).

22. K. Abe, H. Takahashi, H. Suge, Localization of Cells Containing Sedimented Amyloplasts in the Shoots of Normal and Lazy Rice Seedlings. Biol. Sci. Space 8, 221–225 (1994).

23. H. Watanabe, M. Saigusa, S. Morita, Identification of Casparian Bands in the Mesocotyl and Lower Internodes of Rice (*Oryza sativa* L.) Seedlings Using Fluorescence Microscopy. Plant Production Science 9, 390–394 (2006).

24. W. Wang, H. Gao, Y. Liang, J. Li, Y. Wang, Molecular basis underlying rice tiller angle: Current progress and future perspectives. Molecular Plant 15, 125–137 (2022).

25. L. Zhong, L. Geng, Y. Xiang, X. Guang, L. Cao, J. Shi, W. Li, J. Wang, W. He, L. Huang, F. Yang, Y.-X. Bai, S. K. Sahu, X. Guo, S. Zhang, G. Zhang, X. Xu, F. Hu, W. Yang, H. Liu, Y. Zhao, J. Lyu, Comparative spatial transcriptomics reveals root dryland adaptation mechanism in rice and HMGB1 as a key regulator. Molecular Plant 18, 797–819 (2025).

26. L. Abas, M. Kolb, J. Stadlmann, D. P. Janacek, K. Lukic, C. Schwechheimer, L. A. Sazanov, L. Mach, J. Friml, U. Z. Hammes, Naphthylphthalamic acid associates with and inhibits PIN auxin transporters. Proc. Natl. Acad. Sci. U.S.A. 118, e2020857118 (2021).

27. N. Zhang, H. Yu, H. Yu, Y. Cai, L. Huang, C. Xu, G. Xiong, X. Meng, J. Wang, H. Chen, G. Liu, Y. Jing, Y. Yuan, Y. Liang, S. Li, S. M. Smith, J. Li, Y. Wang, A Core Regulatory Pathway Controlling Rice Tiller Angle Mediated by the *LAZY1* - Dependent Asymmetric Distribution of Auxin. Plant Cell 30, 1461–1475 (2018).

28. J.-I. Itoh, K.-I. Nonomura, K. Ikeda, S. Yamaki, Y. Inukai, H. Yamagishi, H. Kitano, Y. Nagato, Rice Plant Development: from Zygote to Spikelet. Plant and Cell Physiology 46, 23–47 (2005).

29. J. Friml, E. Benková, I. Blilou, J. Wisniewska, T. Hamann, K. Ljung, S. Woody, G. Sandberg, B. Scheres, G. Jürgens, K. Palme, AtPIN4 Mediates Sink-Driven Auxin Gradients and Root Patterning in Arabidopsis. Cell 108, 661–673 (2002).

30. T. Garg, Z. Singh, K. Chennakesavulu, K. K. K. Mushahary, A. K. Dwivedi, V. Varapparambathu, H. Singh, R. S. Singh, D. Sircar, D. Chandran, K. Prasad, M. Jain, S. R. Yadav, Species-specific function of conserved regulators in orchestrating rice root architecture. Development, dev.200381 (2022).

31. G. Brunoud, D. M. Wells, M. Oliva, A. Larrieu, V. Mirabet, A. H. Burrow, T. Beeckman, S. Kepinski, J. Traas, M. J. Bennett, T. Vernoux, A novel sensor to map auxin response and distribution at high spatio-temporal resolution. Nature 482, 103–106 (2012).

32. P. Barbier De Reuille, A.-L. Routier-Kierzkowska, D. Kierzkowski, G. W. Bassel, T. Schüpbach, G. Tauriello, N. Bajpai, S. Strauss, A. Weber, A. Kiss, A. Burian, H. Hofhuis, A. Sapala, M. Lipowczan, M. B. Heimlicher, S. Robinson, E. M. Bayer, K. Basler, P. Koumoutsakos, A. H. Roeder, T. Aegerter-Wilmsen, N. Nakayama, M. Tsiantis, A. Hay, D. Kwiatkowska, I. Xenarios, C. Kuhlemeier, R. S. Smith, MorphoGraphX: A platform for quantifying morphogenesis in 4D. eLife 4, e05864 (2015).

33. H. Rakusová, J. Gallego-Bartolomé, M. Vanstraelen, H. S. Robert, D. Alabadí, M. A. Blázquez, E. Benková, J. Friml, Polarization of PIN3-dependent auxin transport for hypocotyl gravitropic response in *Arabidopsis thaliana*. The Plant Journal 67, 817–826 (2011).

34. H. Rakusová, M. Abbas, H. Han, S. Song, H. S. Robert, J. Friml, Termination of Shoot Gravitropic Responses by Auxin Feedback on PIN3 Polarity. Current Biology 26, 3026–3032 (2016).

35. Y. Coudert, Thi, Van Anh Le, P. Gantet, “Rice: A Model Plant to Decipher the Hidden Origin of Adventitious Roots” in Plant Roots: The Hidden Half, A. Eshel, T. Beeckman, Eds. (CRC Press, ed. 4, 2013; https://www.taylorfrancis.com/books/9781439846490/chapters/10.1201/b14550-15), pp. 164–173.

36. Y. Liang, Y. Yuan, Y. Cai, M. Ma, X. Lin, X. Sun, H. Zhao, J. Zhai, N. Han, J. Zhou, W. Wang, Y. Wang, N. Zhang, HSFA2D – LAZY6–LAZY1 module regulates shoot gravitropism and tiller angle in rice. New Phytologist 247, 625–636 (2025).

37. Y. Inukai, T. Sakamoto, M. Ueguchi-Tanaka, Y. Shibata, K. Gomi, I. Umemura, Y. Hasegawa, M. Ashikari, H. Kitano, M. Matsuoka, Crown rootless1, which is essential for crown root formation in rice, is a target of an AUXIN RESPONSE FACTOR in auxin signaling. Plant Cell 17, 1387–1396 (2005).

38. X. Wang, X. Cai, H. Huo, H. Chen, Gravity sensing in plants. New Phytologist, nph.71324 (2026).

39. N. Kawamoto, M. T. Morita, Gravity sensing and responses in the coordination of the shoot gravitropic setpoint angle. New Phytologist 236, 1637–1654 (2022).

40. Y. Song, G. Li, J. Nowak, X. Zhang, D. Xu, X. Yang, G. Huang, W. Liang, L. Yang, C. Wang, V. Bulone, Z. Nikoloski, J. Hu, S. Persson, D. Zhang, The Rice Actin-Binding Protein RMD Regulates Light-Dependent Shoot Gravitropism. Plant Physiol. 181, 630–644 (2019).

41. M. T. Morita, Directional Gravity Sensing in Gravitropism. Annu. Rev. Plant Biol. 61, 705–720 (2010).

42. T. T. Xiao, R. Van Velzen, O. Kulikova, C. Franken, T. Bisseling, Lateral root formation involving cell division in both pericycle, cortex and endodermis is a common and ancestral trait in seed plants. Development, dev.182592 (2019).

43. H. Singh, Z. Singh, R. Kashyap, S. R. Yadav, Lateral root branching: evolutionary innovations and mechanistic divergence in land plants. New Phytologist 238, 1379–1385 (2023).

44. Y. Cai, L. Huang, Y. Song, Y. Yuan, S. Xu, X. Wang, Y. Liang, J. Zhou, G. Liu, J. Li, W. Wang, Y. Wang, LAZY3 interacts with LAZY2 to regulate tiller angle by modulating shoot gravity perception in rice. Plant Biotechnology Journal 21, 1217–1228 (2023).

45. W. Wang, L. Huang, Y. Song, S. Gui, J. Cao, H. Zhang, M. Du, J. Chen, Z. Wang, J. Zhou, X. Meng, D. Zeng, J. Li, Y. Wang, LAZY4 acts additively with the starch–statolith-dependent gravity-sensing pathway to regulate shoot gravitropism and tiller angle in rice. Plant Communications 5, 100943 (2024).

46. E. K. Yoon, S. Dhar, M.-H. Lee, J. H. Song, S. A. Lee, G. Kim, S. Jang, J. W. Choi, J.-E. Choe, J. H. Kim, M. M. Lee, J. Lim, Conservation and Diversification of the SHR-SCR-SCL23 Regulatory Network in the Development of the Functional Endodermis in Arabidopsis Shoots. Molecular Plant 9, 1197–1209 (2016).

47. C. Ortiz-Ramírez, B. Guillotin, X. Xu, R. Rahni, S. Zhang, Z. Yan, P. Coqueiro Dias Araujo, E. Demesa-Arevalo, L. Lee, J. Van Eck, T. R. Gingeras, D. Jackson, K. L. Gallagher, K. D. Birnbaum, Ground tissue circuitry regulates organ complexity in maize and *Setaria*. Science 374, 1247–1252 (2021).

48. M. R. Duvall, G. H. Learn, L. E. Eguiarte, M. T. Clegg, Phylogenetic analysis of rbcL sequences identifies Acorus calamus as the primal extant monocotyledon. Proc. Natl. Acad. Sci. U.S.A. 90, 4641–4644 (1993).

49. M. W. Chase, Monocot relationships: an overview. American J of Botany 91, 1645–1655 (2004).

50. T. Shi, C. Huneau, Y. Zhang, Y. Li, J. Chen, J. Salse, Q. Wang, The slow-evolving Acorus tatarinowii genome sheds light on ancestral monocot evolution. Nat. Plants 8, 764–777 (2022).

51. G. Tian, L. Gao, Y. Kong, X. Hu, K. Xie, R. Zhang, N. Ling, Q. Shen, S. Guo, Improving rice population productivity by reducing nitrogen rate and increasing plant density. PLoS ONE 12, e0182310 (2017).

52. K. Xie, B. Minkenberg, Y. Yang, Boosting CRISPR/Cas9 multiplex editing capability with the endogenous tRNA-processing system. Proc. Natl. Acad. Sci. U.S.A. 112, 3570–3575 (2015).

53. S. Toki, N. Hara, K. Ono, H. Onodera, A. Tagiri, S. Oka, H. Tanaka, Early infection of scutellum tissue with *Agrobacterium* allows high-speed transformation of rice. The Plant Journal 47, 969–976 (2006).

54. D. Kurihara, Y. Mizuta, Y. Sato, T. Higashiyama, ClearSee: a rapid optical clearing reagent for whole-plant fluorescence imaging. Development 142, 4168– 4179 (2015).

55. R. Ursache, T. G. Andersen, P. Marhavý, N. Geldner, A protocol for combining fluorescent proteins with histological stains for diverse cell wall components. The Plant Journal 93, 399–412 (2018).

56. J. Schindelin, I. Arganda-Carreras, E. Frise, V. Kaynig, M. Longair, T. Pietzsch, S. Preibisch, C. Rueden, S. Saalfeld, B. Schmid, J.-Y. Tinevez, D. J. White, V. Hartenstein, K. Eliceiri, P. Tomancak, A. Cardona, Fiji: an open-source platform for biological-image analysis. Nat Methods 9, 676–682 (2012).

57. Z. Xu, W. Wang, T. Yang, L. Li, X. Ma, J. Chen, J. Wang, Y. Huang, J. Gould, H. Lu, W. Du, S. K. Sahu, F. Yang, Z. Li, Q. Hu, C. Hua, S. Hu, Y. Liu, J. Cai, L. You, Y. Zhang, Y. Li, W. Zeng, A. Chen, B. Wang, L. Liu, F. Chen, K. Ma, X. Xu, X. Wei, STOmicsDB: a comprehensive database for spatial transcriptomics data sharing, analysis and visualization. Nucleic Acids Research 52, D1053–D1061 (2024).

