## Supplementary Information for "Spatial Decoupling of the Endodermal Functions Establishes the Innermost Cortex as the Gravity Sensor in Rice Stems"

**Supplementary Informations:** Figures S1 to S12, Tables S1 and S2, and SM References

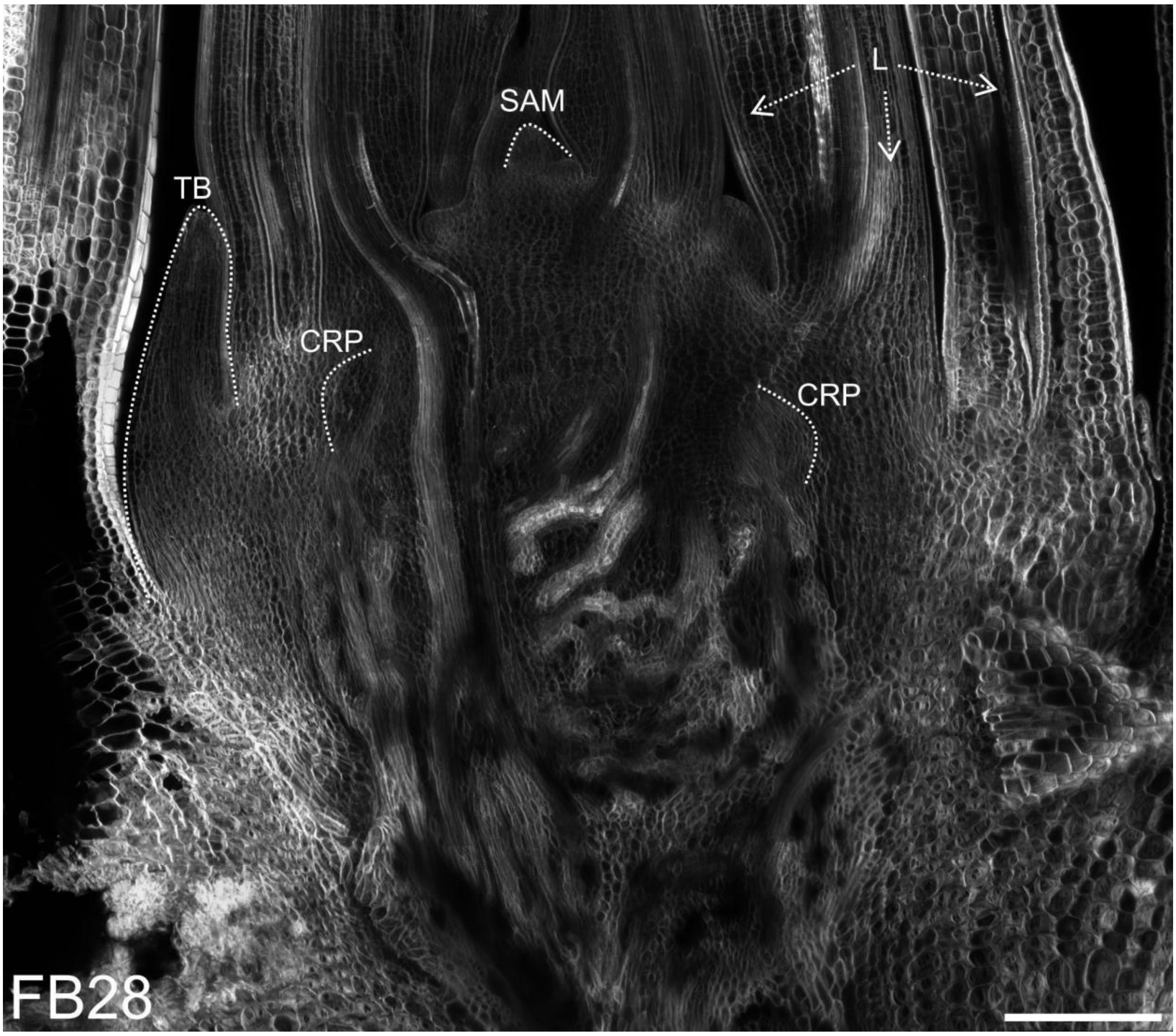


**Fig. S1.** The rice stem produces crown roots as crown root primordia (CRPs), tiller buds (TB), and leaves (L) by the activity of the shoot apical meristem (SAM), making the post-embryonic organs of the rice plant (Scale bar: 200 µm).

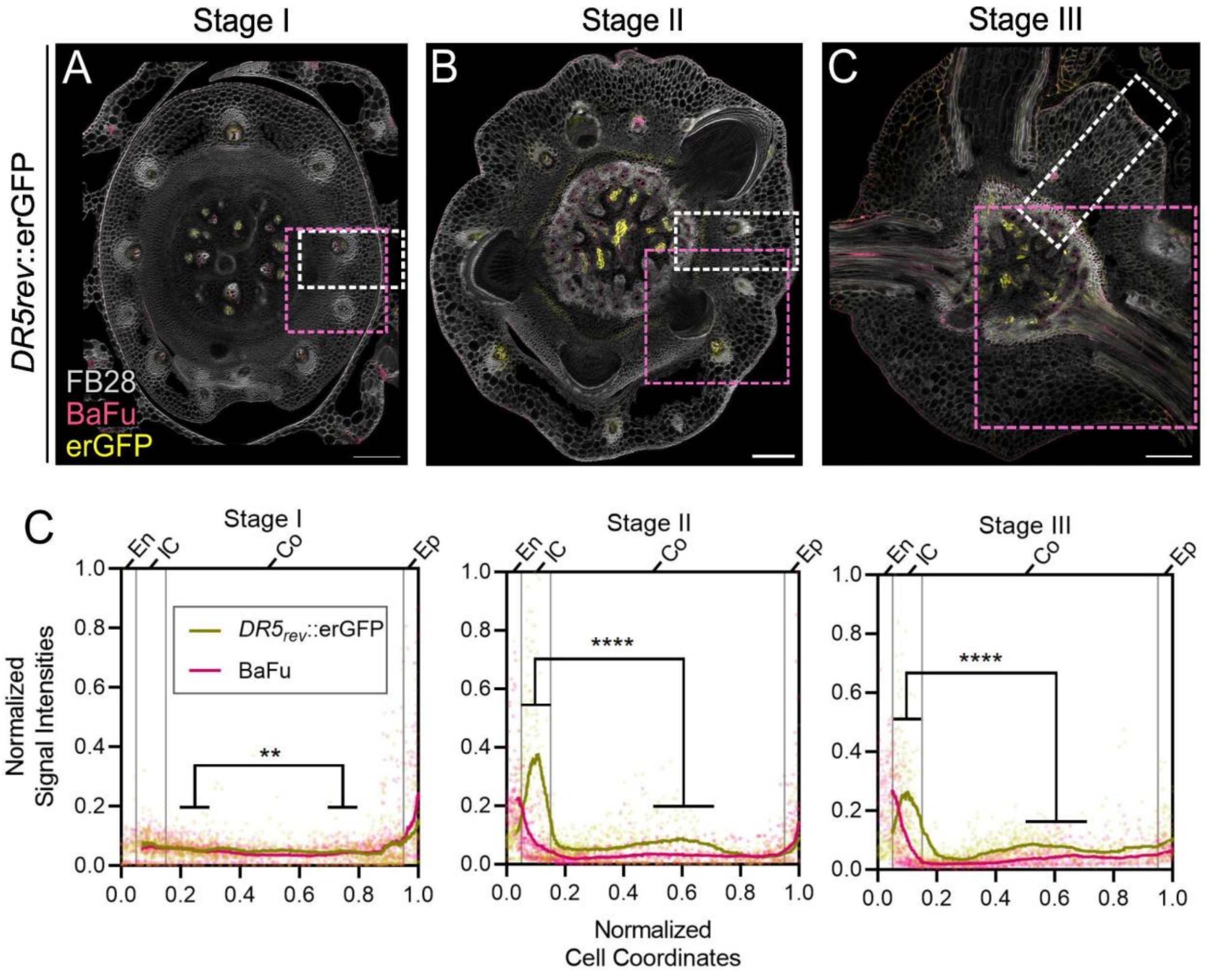


**Fig. S2. Localized auxin response maxima in the innermost cortical cell layer (IC) coincide with En lignification.** *(A–C)*, Representative sections of 6-day-old stem bases across the three stages, Stage I *(A)*, II *(B)*, and III *(C)*, stained with Fluorescent Brightener (FB28) and Basic Fuchsin (BaFu). The dotted white rectangles mark the segmentation sectors, and the dotted pink squares mark the cropped image used for Fig. 2*A–C* (Scale bars: 100 µm). *(D)*, Graphical representation of normalized BaFu and *DR5rev*::erGFP signal intensities, plotted against normalized cell coordinates (0 = endodermis and 1 = epidermis), across the three stages (n = 6 sections from 6 independent seedlings for each stage). ****$p < 0.0001$ and **$p = 0.0052$ from Mann-Whitney U test on *DR5rev*::erGFP signal intensities. Data from the same segmentations as for Fig. 2 *B–C*. Note the peak of the *DR5* signal at Stage II, adjacent to the BaFu signal coming from the lignified endodermis. Each data point on the plot represents an individual cell, with solid lines depicting the rolling mean.

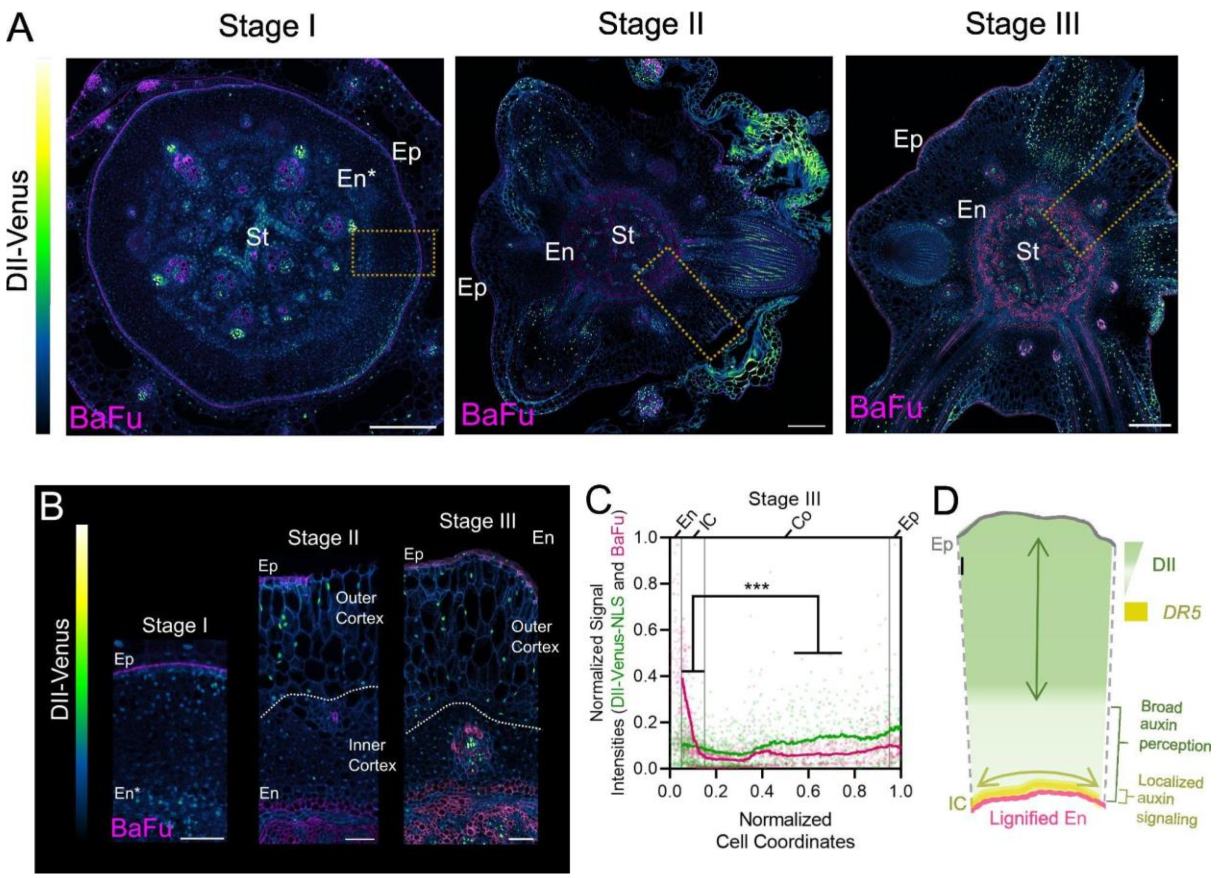


**Fig. S3. The inner cortex shows broad auxin perception, but selective transcriptional activation of auxin signaling in the IC.** *(A)*, Representative vibratome sections of DII-Venus lines stained with Basic Fuchsin (BaFu) across the three stages of stem development (Stage I, II, and III). Dotted orange rectangles mark the cropped sectors of B (Scale bar: 200 µm). *(B)*, Cropped orange-dotted sectors from *(A)* between endodermis (En, En* marks the unlignified endodermis in Stage I) and epidermis (Ep). Note the nuclear signal intensities at Stage I, which are uniform throughout the cortex. At Stages II and III, the intensities remain lower in the inner cortex and increase in the outer cortex (separated by the dotted white line, Scale bar: 20 µm). *(C)*, Graphical representation of normalized DII-Venus and BaFu signal intensities plotted against normalized cell coordinates at Stage III (0 = endodermis and 1 = epidermis, n = 6, $p^{***} = 0.0002$ from Mann-Whitney U test). *(D)*, A schematic summary showing the broad perception of auxin (DII) but a localized signaling output (*DR5*) that coincides with the appressed orientation of cells in the IC, adjacent to the lignified En.

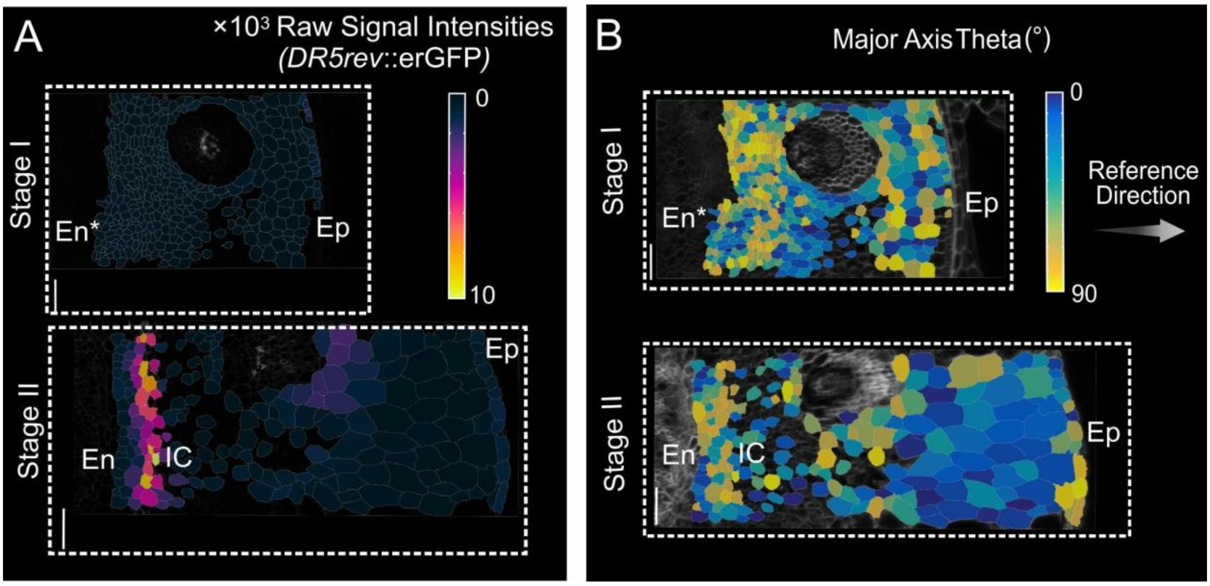


**Fig. S4. Cellular segmentations of the rice stem bases at Stage I and II** (dashed white rectangles from Fig. S2 *A* and *B*) showing raw *DR5rev*::erGFP signal intensities *(A)* and the Major Axis Theta *(B)* (MAθ, angle in degrees between the Major Axis of the cell and the reference direction; En: Endodermis, Ep: Epidermis; IC: Innermost Cortical Cell Layer, Scale bar: 50 µm).

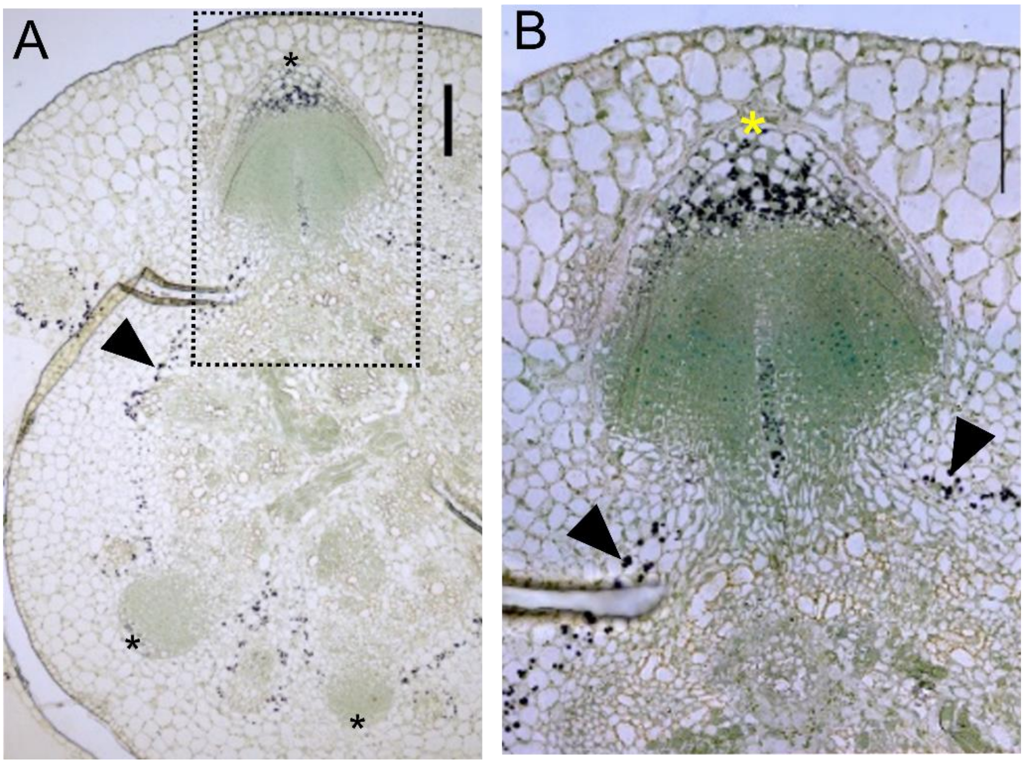


**Fig. S5. The IC shows statolith accumulation.** *(A)*, Lugol-stained section of a rice stem base showing a ring of statoliths (black arrowhead) around the CRPs in the innermost layer of the cortex (black asterisks) (Scale bar: 100 μm). ***(B)*,** Cropped section from A, showing the statolith-rich IC (black arrowheads), and statoliths in the developing columella of the CRP (yellow asterisk) (Scale bar: 100 μm).

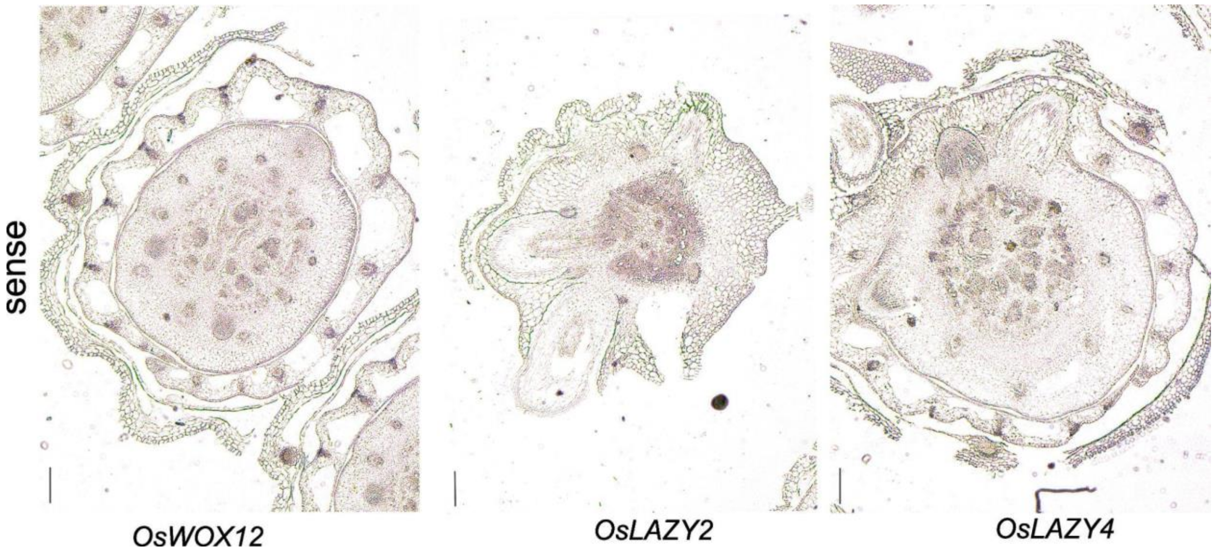


**Fig. S6. Sense probe controls for *OsWOX12, OsLAZY2,* and *OsLAZY4.*** Cross-section of rice stem base probed with sense riboprobes of *OsWOX12*, *OsLAZY2* and *OsLAZY4* (Scale bar: 200 μm).

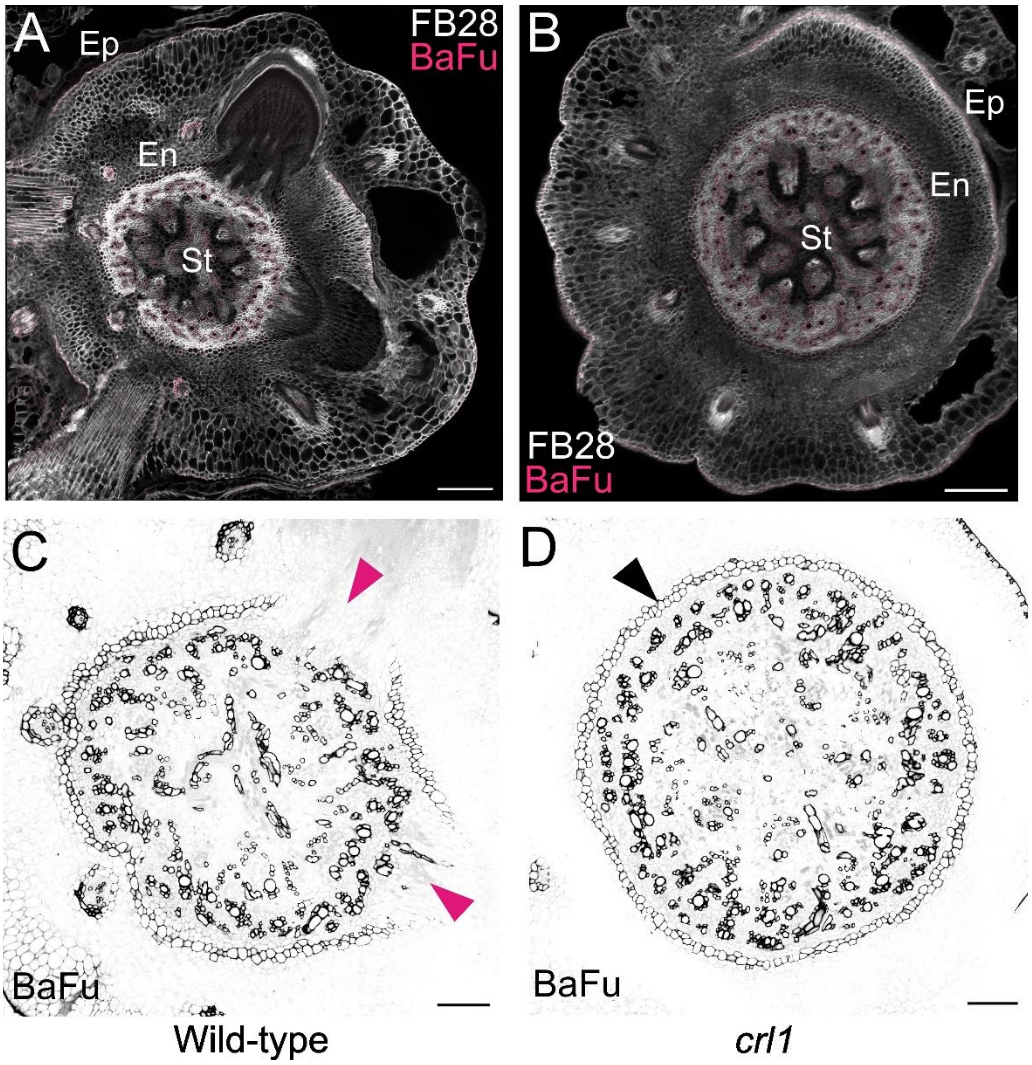


**Fig. S7. *crl1* shows a continuous lignified endodermis.** *(A-B)*, Basic Fuchsin (BaFu) and Fluorescent Brightener 28 (FB28) stained sections of wild-type (Taichung 65, *A*) and *crl1* (*B*) (Scale bar: 200 µm; Ep: Epidermis, En: Endodermis, and St: Stele). *(B-C)*, Contrast-enhanced BaFu channel from *A* and *B*, note the continuous En (black arrowhead) in *crl1* (*D*), while WT (*C*) shows discontinuity at the site of origin of CRPs (pink arrowheads) (Scale bar: 100 µm).

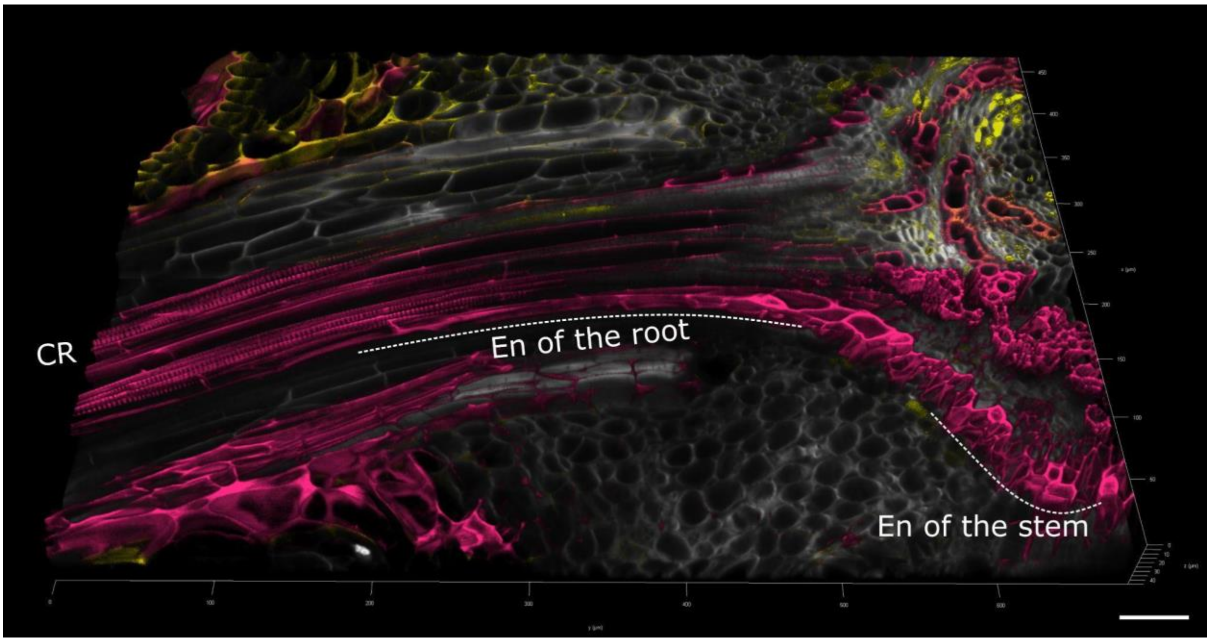


**Fig. S8. 3D reconstruction of a section showing that the endodermis of the stem is continuous with that of the endodermis of the crown root (CR).** This ensures the continuity of the apoplastic barrier among organs (Scale bar: 100 µm).

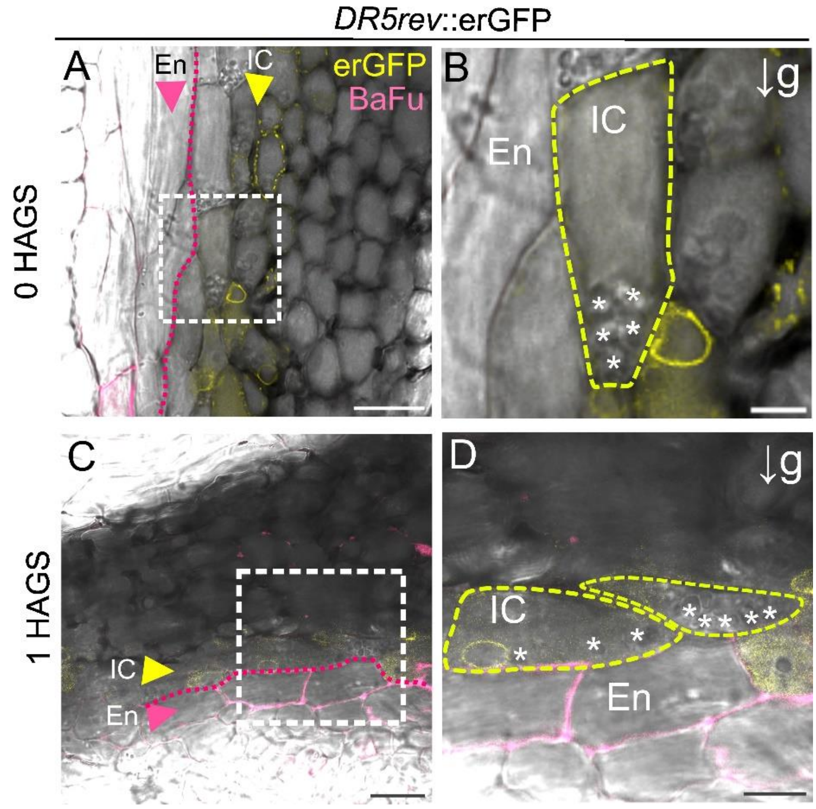


**Fig. S9. Sedimentation of amyloplasts in the IC upon gravistimulation.** (*A*), En and the IC as visible under brightfield at 0 HAGS (Scale bar: 20 µm). (*B*), White-dashed square from (*A*), the yellow-dashed line marks the outline of one of the cells of the IC, note the amyloplasts (white asterisks) sedimented along the direction of the gravity (Scale bar: 5 µm). (*C*), En and IC 1 HAGS (Scale bar: 50 µm). (*D*), White-dashed square from (*C*), the yellow-dashed line marks the outline of cells of the IC, note the amyloplasts (white asterisks) sedimented along the direction of the gravity (Scale bar: 10 µm).

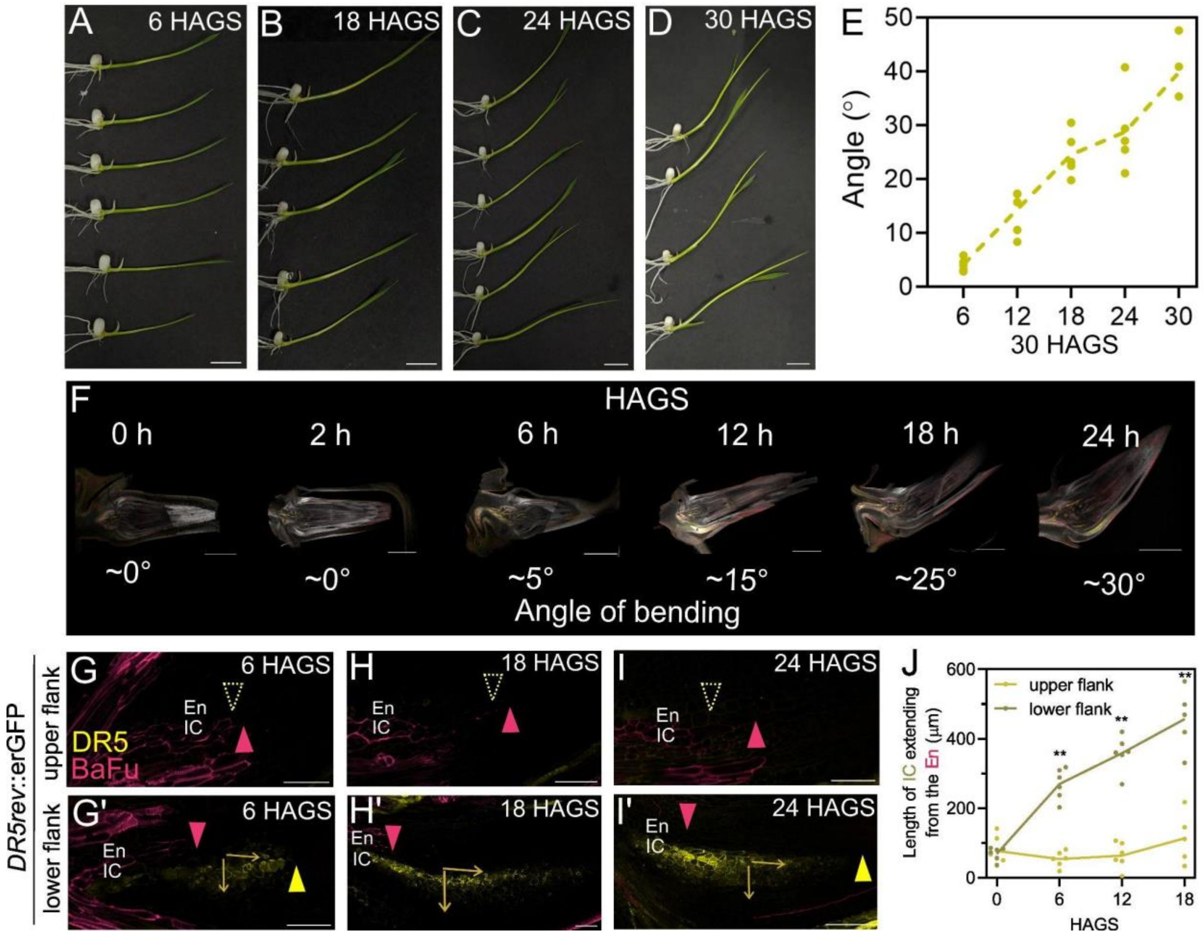


**Fig. S10. 90° gravistimulation experiment with *DR5rev*::erGFP lines.** (*A*-*D*) Seedlings hours after gravistimulation (HAGS, Scale bar: 1 cm). *(E)*, Angle of bending (°) HAGS, average angles from this were used for rotating sections in Fig. S8*F*. *(F)*, Gravistimulation experiment on *DR5rev*::erGFP lines showing representative longitudinal sections from 0 to 24 HAGS (Scale bar: 1 mm). *(G-I')*, Upper (*G*-*I*) and lower flanks (*G'*-*I'*) of sections from Fig. S5*F* at 6 (*G* and *G'*), 18 (*H* and *H'*), 24 (*I* and *I'*) HAGS. The yellow and pink arrowheads mark the positions of lignified En and the *DR5* signal of the IC. Note the disappearing signal in the upper flanks, denoted by dashed-yellow arrowheads (Scale bar: 50 µm for each. For *I'*: 100 µm). *(J)*, Quantitative representation of the disparity between the lignified En and the extending IC signal in the lower vs upper flanks (n=6 for 0, 6, and 12 HAGS; n=5 for 18 HAGS. $^{**}p = 0.0022$ for 6 and 12 HAGS, $^{**}p = 0.0079$ for 18 HAGS, and $p = 0.82$ for 0 HAGS from Mann-Whitney U test).

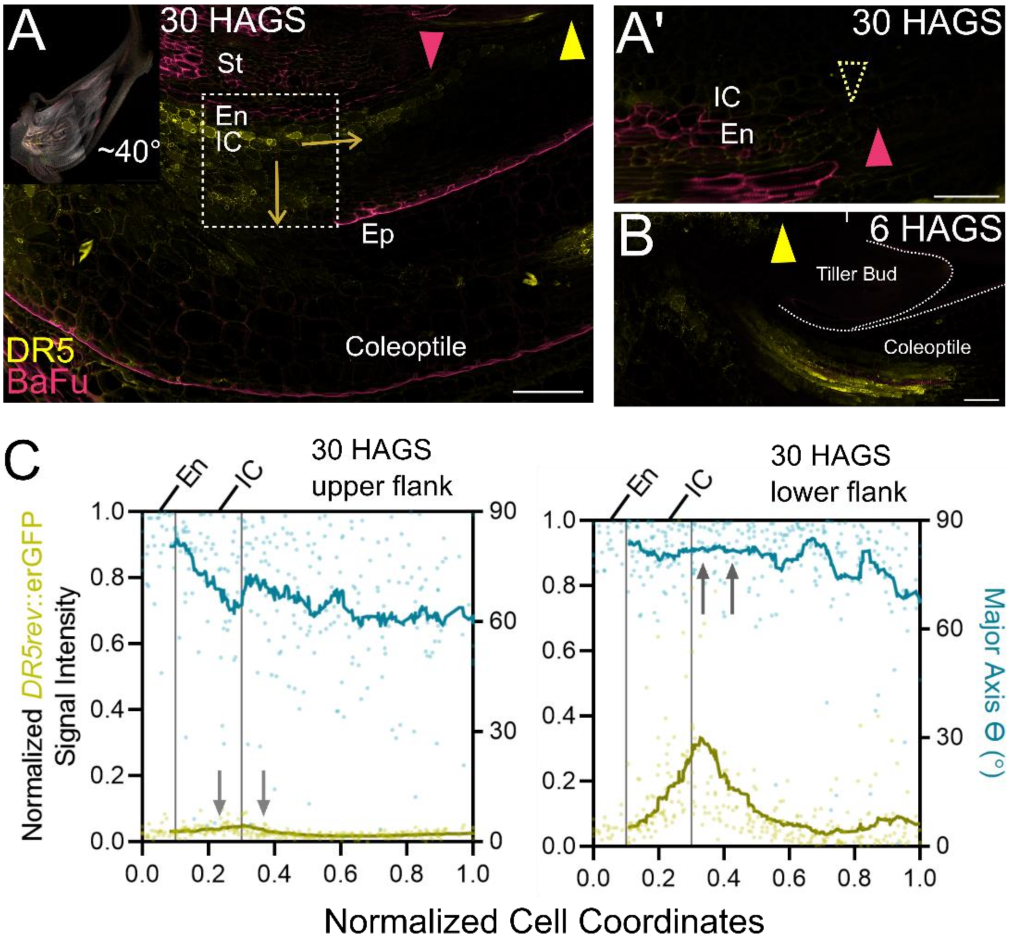


**Fig. S11. The lower flank shows an expanding *DR5* signal and circumferential anisotropy.** *(A),* Lower flank at 30 h after gravistimulation (30 HAGS) of the inset longitudinal section. The dotted square marks the region used for quantification in MorphoGraphX (Scale bar of inset: 1 mm. Scale bar: 100 µm). *(A')* The corresponding upper flank at 30 HAGS (Scale bar: 50 µm). *(B), DR5* signal in the coleoptile of the stem, visible at 6 HAGS (Scale bar: 50 µm). *(C),* Graphical representation of normalized *DR5rev*::erGFP signal intensities and Major Axis Theta (MAθ in °), plotted against normalized cell coordinates (0 = endodermis and 1 = stem coleoptile junction). The *DR5* signal was normalized to the range between the highest and lowest values within the lower and upper flanks. Note the reduced *DR5* signal in the upper flank and the increased MAθ in the lower flank (grey arrows).

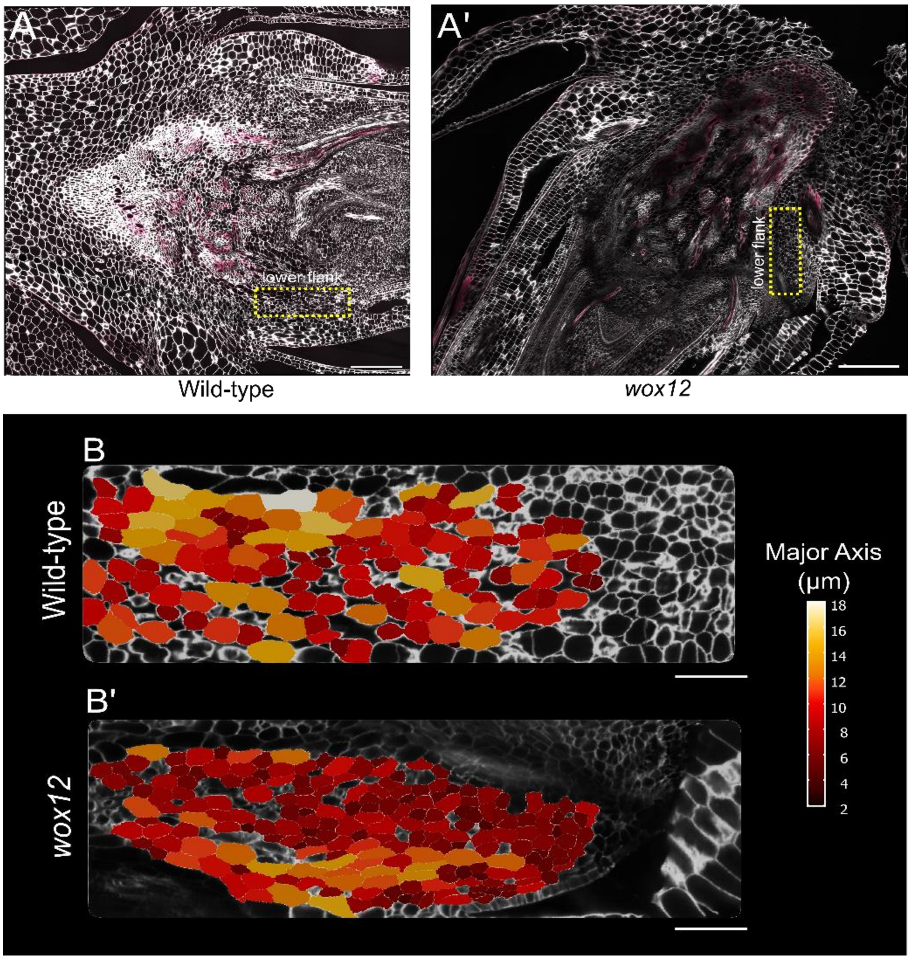


**Fig S12. The *wox12* mutant shows reduced cell elongation in the lower flank after gravistimulation.** *(A-A'),* Representative longitudinal vibratome sections of wild-type *(A)* and *wox12 (A').* The dashed-yellow rectangle marks the uniform region (450 µm × 150 µm) used for the segmentations (Scale bar: 200 µm). *(B-B'),* Representative MorphoGraphX segmentations at the lower flank of the wild-type *(B)*, and *wox12 (B'),* the heatmap shows the major axis (µm) (Scale bar: 50 µm).

**Table S1.** Genes and their RAP DB IDs used to generate Fig. 3D and E. *Log*$_2$_FC (fold-change) and FDR were calculated from CPM values for Bin 1-2 *vs*. Bin 3-10.

| **RAP DB ID** | **Common Name** | **Role** | ***Log*$_2$_FC (Bins 1-2 vs. Bin 3-10)** | **FDR** | **Reference** |
|---|---|---|---|---|---|
| Os06g0166500 | *OsIAA20* | Auxin redistribution | 2.750 | 0.001 | *(1)* |
| Os03g0325600 | *OsWOX12* | Asymmetric growth | 2.711 | 0.006 | *(2)* |
| Os02g0228900 | *OsIAA7* | Auxin redistribution | 1.621 | 0.001 | *(3)* |
| Os02g0743400 | *OsPIN1b* | Auxin redistribution | 1.512 | 0.012 | *(4)* |
| Os06g0232300 | *OsPIN1a* | Auxin redistribution | 1.381 | 0.006 | *(4)* |
| Os03g0735000 | *OsAGPL1* | Gravity sensing | 1.341 | 0.142 | *(5)* |
| Os06g0677800 | *OsARF17* | Auxin redistribution | 1.210 | 0.001 | *(6)* |
| Os12g0601300 | *OsIAA30* | Auxin redistribution | 1.011 | 0.002 | *(3)* |
| Os05g0421300 | *OsLAZY6* | Auxin redistribution | 1.008 | 0.019 | *(7)* |
| Os12g0613700 | *OsARF25* | Auxin redistribution | 0.943 | 0.018 | *(6)* |
| Os04g0671900 | *OsARF12* | Auxin redistribution | 0.920 | 0.016 | *(6)* |
| Os02g0450000 | *OsTAC4* | Auxin redistribution | 0.898 | 0.128 | *(8)* |
| Os02g0141100 | *OsARF5* | Auxin redistribution | 0.883 | 0.112 | *(3)* |
| Os02g0180200 | *OsLAZY2* | Gravity sensing | 0.813 | 0.142 | *(9)* |
| Os03g0161900 | *OsHSFA2D* | Auxin redistribution | 0.538 | 0.552 | *(7)* |
| Os03g0133300 | *OsLAZY3* | Gravity sensing | 0.147 | 0.754 | *(10)* |
| Os05g0408800 | *OsLAZY4* | Gravity sensing | -0.419 | 1.000 | *(11, 12)* |
| Os11g0490600 | *OsLAZY1* | Auxin redistribution | -0.837 | 0.272 | *(13)* |
| Os03g0237250 | *OsLPA1* | Gravity sensing | -0.945 | 1.000 | *(14)* |
| Os07g0684900 | *OsWOX11* | Asymmetric growth | -14.535 | 0.509 | *(2)* |

**Table S2.** Primers used in the study for cloning probes for RNA *in situ* hybridization.

| RAP DB ID | Common Name | Primer Sequence (5'-3') | |
|---|---|---|---|
| Os02g0180200 | *OsLAZY2* | FP | CTTAGGAATGCCAGCAGGCCTT |
| | | RP | CAGCTAACAGTACTCGCCACG |
| Os05g0408800 | *OsLAZY4* | FP | ACTCGCTCCCTCCTCCAAAG |
| | | RP | GCGGCCATTAGGGTTTCGGA |
| Os03g0325600 | *OsWOX12* | FP | CCAAATCGGAGAGAGCTACTTTC |
| | | RP | GCATGCCTAAAAATGTGGTGC |

### SM References


1. H. Zhang, X. Li, D. Sang, L. Huang, Y. Song, M. Du, J. Cao, W. Wang, PROG1 acts upstream of LAZY1 to regulate rice tiller angle as a repressor. *The Crop Journal* **11**, 386–393 (2023).

2. N. Zhang, H. Yu, H. Yu, Y. Cai, L. Huang, C. Xu, G. Xiong, X. Meng, J. Wang, H. Chen, G. Liu, Y. Jing, Y. Yuan, Y. Liang, S. Li, S. M. Smith, J. Li, Y. Wang, A Core Regulatory Pathway Controlling Rice Tiller Angle Mediated by the *LAZY1* -Dependent Asymmetric Distribution of Auxin. *Plant Cell* **30**, 1461–1475 (2018).

3. Y. Hu, D. Xue, S. Wang, Q. Zhang, X. Zhang, J. Yang, Y. Lv, B. Yan, Y. Yin, Z. Cui, T. Li, W. Chen, X. Wang, An auxin response factor regulates tiller angle and shoot gravitropism by directly activating related gene expression in rice. *Journal of Advanced Research* **78**, 147–162 (2025).

4. Y. Li, J. Zhu, L. Wu, Y. Shao, Y. Wu, C. Mao, Functional Divergence of PIN1 Paralogous Genes in Rice. *Plant and Cell Physiology* **60**, 2720–2732 (2019).

5. M. Okamura, T. Hirose, Y. Hashida, T. Yamagishi, R. Ohsugi, N. Aoki, Starch reduction in rice stems due to a lack of OsAGPL1 or OsAPL3 decreases grain yield under low

irradiance during ripening and modifies plant architecture. *Functional Plant Biology* **40**, 1137–1146 (2013).

6. Y. Li, J. Li, Z. Chen, Y. Wei, Y. Qi, C. Wu, OsmiR167a-targeted auxin response factors modulate tiller angle via fine-tuning auxin distribution in rice. *Plant Biotechnol J* **18**, 2015–2026 (2020).

7. Y. Liang, Y. Yuan, Y. Cai, M. Ma, X. Lin, X. Sun, H. Zhao, J. Zhai, N. Han, J. Zhou, W. Wang, Y. Wang, N. Zhang, HSFA2D – LAZY6–LAZY1 module regulates shoot gravitropism and tiller angle in rice. *New Phytologist* **247**, 625–636 (2025).

8. H. Li, H. Sun, J. Jiang, X. Sun, L. Tan, C. Sun, *TAC4* controls tiller angle by regulating the endogenous auxin content and distribution in rice. *Plant Biotechnology Journal* **19**, 64–73 (2021).

9. L. Huang, W. Wang, N. Zhang, Y. Cai, Y. Liang, X. Meng, Y. Yuan, J. Li, D. Wu, Y. Wang, *LAZY2* controls rice tiller angle through regulating starch biosynthesis in gravity-sensing cells. *New Phytologist* **231**, 1073–1087 (2021).

10. Y. Cai, L. Huang, Y. Song, Y. Yuan, S. Xu, X. Wang, Y. Liang, J. Zhou, G. Liu, J. Li, W. Wang, Y. Wang, LAZY3 interacts with LAZY2 to regulate tiller angle by modulating shoot gravity perception in rice. *Plant Biotechnology Journal* **21**, 1217–1228 (2023).

11. W. Wang, L. Huang, Y. Song, S. Gui, J. Cao, H. Zhang, M. Du, J. Chen, Z. Wang, J. Zhou, X. Meng, D. Zeng, J. Li, Y. Wang, LAZY4 acts additively with the starch–statolith-dependent gravity-sensing pathway to regulate shoot gravitropism and tiller angle in rice. *Plant Communications* **5**, 100943 (2024).

12. J. Fan, X. Ma, J. Zou, S. Li, Y. Liu, D. Guo, W. Jiang, X. Li, F. Liu, L. Tan, LATA1 , a RING E3 ligase, modulates the tiller angle by affecting auxin asymmetric distribution and content in rice. *The Plant Journal* **120**, 429–444 (2024).

13. P. Li, Y. Wang, Q. Qian, Z. Fu, M. Wang, D. Zeng, B. Li, X. Wang, J. Li, LAZY1 controls rice shoot gravitropism through regulating polar auxin transport. *Cell Res* **17**, 402–410 (2007).

14. X. Wu, D. Tang, M. Li, K. Wang, Z. Cheng, Loose Plant Architecture1, an INDETERMINATE DOMAIN Protein Involved in Shoot Gravitropism, Regulates Plant Architecture in Rice. *Plant Physiology* **161**, 317–329 (2012).
